# Defense against parasites covaries with reproductive timing, not with resistance

**DOI:** 10.64898/2026.02.28.708748

**Authors:** Amanda K Gibson, Linyao Peng, Tessa Batterton, Neha Channamraju, Victoria Feist, Sarah Hesse, Anne N Janisch, Hongyi Shui

**Affiliations:** Department of Biology, University of Virginia, Charlottesville, VA, USA; Independent researcher, Charlottesville, VA, USA

## Abstract

Defense is the ability of a host to minimize fitness loss to parasites. It is among the most variable phenotypes in host populations, and this variation facilitates rapid adaptation under parasite-mediated selection. We do not, however, know the underlying host traits that explain this variation in defense against parasites. A common assumption is that the most defended hosts are the most resistant, meaning they limit the establishment and growth of infecting parasites. Under this assumption, resistance traits should evolve readily under parasite selection. Resistance is, however, just one of many strategies hosts use to defend against parasites, and it does not consistently covary with fitness in the presence of parasites. We accordingly asked: which host traits covary with defense against parasites? We use controlled exposures to characterize genetic variation in defense of the nematode *Caenorhabditis elegans* against its natural microsporidian parasites. We report extensive variation in defense among wild strains of *C. elegans*: some strains lost 60% of fecundity under parasite exposure, while others were unaffected. We then tested the covariance of defense with two prominent host traits, resistance and reproductive timing. Our results did not support the hypothesis that resistance covaries with defense: strains with lower parasite burden did not have higher relative fecundity under exposure. Our results instead supported the hypothesis that life history covaries with defense: host strains that reproduced quickly had higher relative fecundity under exposure, consistent with the idea that parasites diminish future reproductive opportunities. The observed variation in defense among host strains indicates significant potential for wild *C. elegans* populations to evolve in response to their natural parasites. Because reproductive timing underpins this variation in defense, parasite-mediated selection could operate directly on host life history traits and should also be highly sensitive to shifts in life history driven by other biotic and abiotic factors.

**AUTHOR SUMMARY:** Some hosts fare much better than others in the face of parasite infection. What traits differentiate defended hosts from undefended hosts? The answer to this question is critical for identifying the strategies that best protect hosts from their parasites. It also allows us to predict and interpret the evolution of host populations over the course of epidemics. To address this question, we surveyed wild strains of a tractable model host, the nematode *Caenorhabditis elegans*, for their response to two species of microsporidian parasites. We found that, on average, parasite exposure substantially impaired the ability of hosts to reproduce. Host strains, however, varied widely: some experienced major losses in fecundity with exposure, while others were highly defended, showing little to no change. We identified reproductive timing as a key trait that differentiated defended hosts from undefended hosts. Our results suggest that reproducing quickly may have been protective, by allowing hosts to make most of their offspring before parasites impaired reproduction. We did not find evidence that resistance was protective: host strains with lower parasite burdens did not reproduce better than those with high parasite burdens. These findings give added weight to life history as a major component of host defense against parasites.

## INTRODUCTION

Many different strategies can increase a host’s defense against parasites. Avoidance reduces a host’s exposure to parasites, resistance limits the establishment and proliferation of infecting parasites, and tolerance curbs the damage inflicted by infecting parasites [1,2]. These strategies result from a variety of mechanisms, from life history to behavior to immunity. We define defense against parasites as the net ability of a host to minimize fitness loss to parasites, and it reflects the holistic actions of these different mechanisms.

Natural host populations typically harbor extensive genetic variation in defense. This variation is evident at the phenotypic and genomic levels [3–9]. Indeed, loci linked to defense are often associated with the most variable regions of genomes [10–12]. This extensive variation in defense indicates that host populations can adapt rapidly to selection by parasites. A central question is: which mechanisms explain why some hosts are so much more defended against parasitism than others?

To answer this question, we must identify which heritable traits covary with fitness in the face of parasites [13,14]. Resistance traits are an obvious answer. Resistance describes any defense mechanism that limits the establishment and proliferation of an infection. It is typically quantified as the inverse of infection burden [2,15]. Resistant hosts by definition have lower burdens of parasites, and they are accordingly expected to have higher fitness. There is ample support for this expectation [e.g., 13,16,17]. For example, resistance under artificial exposure to a trematode parasite correlated positively with the frequency in the wild of clones of the snail *Potamopyrgus antipodarum,* in spite of resistant clones having lower baseline fecundity [18]. However, it is reasonably common to find that the fitness consequences of infection are divorced from parasite burden, such that hosts can be relatively defended in spite of having high parasite loads [15]. For example, Kover and Schaal [14] found that bacterial load did not explain variation in the fitness consequences of infection across accessions of *Arabidopsis thaliana* hosts. These conflicting findings suggest that resistance traits may not always feature as the primary component of defense against parasites.

Hosts have access to many alternatives to resistance. Prominent among these is life history. Parasites are likely to curtail a host’s future reproductive opportunities, either through mortality or declining fecundity, such that in the presence of parasites, hosts achieve relatively higher reproductive success when they reproduce earlier [19–21]. Hosts indeed show evidence of shifting their reproductive schedule in response to infection: *Biomphalaria glabrata* snails increased egg laying immediately after exposure to *Schistosoma mansoni* trematodes, as compared to unexposed controls [22]. House crickets behaved similarly (*Acheta domesticus*) after bacterial infection [23]. Parasites may also favor hosts that constitutively reproduce early: at sites where castrating trematodes are prevalent, snails of the species *Cerithideopsis californica* and *Potamopyrgus antipodarum* matured at a smaller size, consistent with early reproductive maturity [24,25]. Likewise, Rancho Grande harlequin toads (*Atelopus cruciger*) matured at smaller size in populations that have experienced chytrid epidemics [26]. These life history shifts differ from resistance traits in that they confer defense by maintaining host fitness without directly reducing parasite fitness. They could thus potentially serve as a mechanism of tolerance, a defense strategy that limits the harm inflicted by a given burden of parasites [2,15,27]. Life history shifts can confer defense even when they are not the result of direct selection by parasites. For example, annual plant species in the genus *Silene* are much more defended against anther-smut fungus *Microbotryum* than are perennial relatives, but it is unlikely the annual habit evolved in response to parasite selection. Increased parasite defense is more likely a correlated response to selection by other forces operating on life history in the *Silene* genus [28,29]. Similarly, when lines of *Aedes aegypti* mosquitoes were experimentally selected for faster development, they experienced a correlated reduction in the fitness effect of a microsporidian parasite [30]. In summary, prior studies indicate that both resistance and life history traits contribute to maintaining host fitness in the face of parasites. We do not know their relative contributions to defense against parasites – does one covary more strongly with defense than the other?

We set out to address this question in the nematode host *Caenorhabditis elegans* and its natural microsporidian parasites in the genus *Nematocida* [31]*. C. elegans* is a globally-distributed free-living nematode that feeds on bacterial blooms growing on rotting vegetation [32]. Most *C. elegans* individuals are self-fertile hermaphrodites, and distinct selfing lineages persist within metapopulations via dispersal among bacterial patches [33–35]. Nematodes inadvertently ingest spores of *Nematocida* while feeding on bacteria [36]. Several *Nematocida* species, including *N. parisii* and *N. ironsii*, establish infections in the intestinal cells and transmit horizontally via a fecal-oral route [36,37]. *Nematocida* infection strongly reduces host fitness, by reducing lifetime fecundity, suppressing population growth rate, and eliminating the fitness of dispersal stage larvae [36,38]. There is variation among wild *C. elegans* strains in their defense against *Nematocida:* the presence of parasites alters which strains increase in frequency in multi-strain competitions [33,39,40]. We do not know the traits that explain this variation in defense among hosts. Prior studies have characterized resistance against *Nematocida* infection [41–45]. Notably some strains have the capacity to clear *Nematocida* infection, and they accordingly have elevated fitness in the presence of parasites [39,40]. Other host strains, however, appear to maintain high fitness in the presence of *Nematocida* without limiting parasite burden [40], suggesting resistance traits may not fully explain variation in defense. Host strains also vary substantially in life history traits, though whether these contribute to variation in defense against parasites is unknown [46].

In this study, we characterized variation in defense among *C. elegans* strains sampled from the Hawaiian Islands. Populations of *C. elegans* from the Hawaiian Islands harbor substantially more genetic diversity than those from regions outside the Pacific [47,48], and thus Hawaiian strains offer valuable insight on the extent of genetic variation in host traits. We quantified variation in defense by comparing the fecundity of host strains in the presence of parasites, relative to their fecundity in the absence of parasites. We exposed hosts to two parasite species – *N. parisii* and *N. ironsii –* at two doses to gain a general sense of a host strain’s level of defense and to determine the sensitivity of defense to specific exposure conditions. We then tested two hypotheses for the traits that explain the observed variation in defense. First, we quantified variation in life history traits and tested whether host strains that reproduced earlier maintained higher fitness in the face of parasites. Second, we quantified variation in resistance traits and tested whether host strains with lower infection prevalence and load maintained higher fitness in the face of parasites. We found extensive variation in defense against parasites that covaried strongly with the timing of host reproduction.

## RESULTS

To test our hypotheses, we characterized traits of 19 Hawaiian strains that broadly represent the major phylogenetic clusters of Hawaiian *C. elegans*; these clusters are referred to as the Volcano, Divergent, Low, and Invaded groups [47]. We also included the canonical lab strain N2 for reference (S1 Fig, Table A in S1 Text). We first report the fitness consequences of parasite exposure and evaluate the extent of genetic variation in defense against parasites. We then report tests of the covariance of life history and resistance traits with the observed variation in defense.

### Genetic variation in defense against parasites

We characterized variation in defense against *Nematocida* by estimating each host strain’s mean fitness reduction following exposure to parasites. Host fitness was estimated as lifetime fecundity. We paired a standard fecundity assay with a computer vision pipeline to measure host survival and the number of viable eggs produced each day by individual hermaphrodites in the absence (control) and presence (exposed) of parasites. These data provided daily and total offspring production, from which we inferred mean baseline fecundity, fecundity schedule, and defense for host strains. Host strains with the largest mean reductions in fecundity following exposure were considered to have relatively low defense against parasites. To broadly evaluate defense, hosts were exposed to two species of *Nematocida, N. parisii* and *N. ironsii*, at low and high doses (Table B in S1 Text). This gave a total of five treatments, one control and four exposed. In total, we captured survival and reproduction over the course of 144 hours for 1,028 hermaphrodites, with 203-209 per treatment and 38-50 per Hawaiian strain (Table C in S1 Text).

Exposure to parasites reduced total fecundity by an average of 43%, from 148.4 ± 4.9 offspring per host in the control treatment to 84.6 ± 1.9 on average across exposed treatments (mean ± standard error; Fig. 1a, Table D in S1 Text). In the absence of parasites, host strains showed continuous variation in fecundity, ranging from 29.4 ± 12.3 offspring per host for XZ1514 (ID #1) to 226.9 ± 12.5 for N2 (ID #20). We estimated broad sense heritability for baseline fecundity to be 0.39 [95% posterior probability: 0.19, 0.59]. Host strains also showed continuous variation in total fecundity in exposed conditions: averaging over all exposure treatments, fecundity ranged from 9.8 ± 2.8 offspring per host for XZ1514 (#1) to 134.8 ± 9.5 ECA705 (#5), yielding a broad sense heritability estimate of 0.29 [0.15,0.46] (Fig. 1b, Table D in S1 Text). Estimates of total offspring number included hosts that did not reproduce and therefore incorporated variation in host survival in the presence of parasites. We report survival effects in S2 Text.

**Figure 1:**
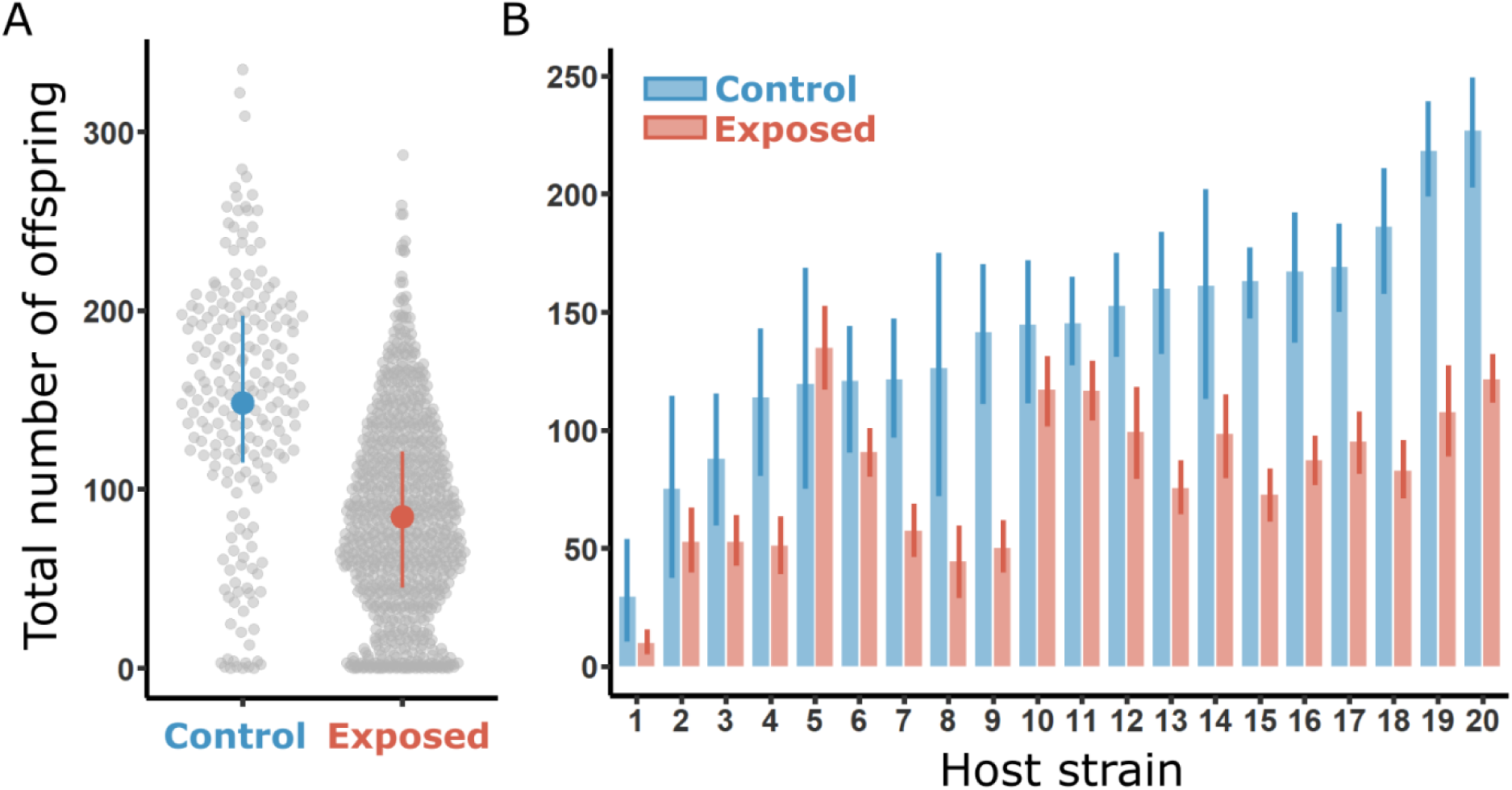
Parasite exposure reduces fecundity. **A** displays the fecundity of hosts in control (blue) and exposed conditions (red). Gray points show the total number of offspring for individual hermaphrodites, across all strains. Filled points indicate the mean per treatment, and error bars show the interquartile ranges of the data. The Control group represents data from 209 hosts across 20 strains, while the Exposed group represents data from 819 hosts from 20 strains and four exposure treatments. **B** shows the fecundity of host strains in control (blue) and exposed conditions (red). Bars show the mean number of offspring per host strain, and error bars show 95% confidence intervals. Host strains are ordered from left to right by increasing fecundity in baseline control conditions. Host strains are indicated by their numeric ID (Table A in S1 Text): 1=XZ1514; 2 = ECA812; 3= ECA740; 4=ECA730; 5=ECA705; 6=ECA724; 7=ECA363; 8=ECA1997; 9=CB4856; 10=ECA372; 11=QX1792; 12=ECA743; 13=ECA2334; 14=DL238; 15=ECA347; 16=ECA1977; 17=ECA746; 18=QX1791; 19=ECA744; 20=N2

We quantified defense against parasites as average fecundity under parasite exposure relative to average baseline fecundity in the control treatment. Host strains that maintained a greater proportion of their baseline fecundity following exposure were considered more defended. Host strains showed continuous variation in their level of defense (Fig. 2a; Table D in S1 Text – likelihood ratio test of the interaction of host strain and exposure: *χ^2^*=74.6, df=19, p<0.001). Several strains were poorly defended, losing on average ∼60% or more of their total fecundity with parasite exposure (i.e., #1- XZ1514, #8 - ECA1997, #9 - CB4856, #18 - QX1791). Other strains were consistently highly defended: on average, ECA724 (#6), QX1792 (#11), and ECA372 (#10) lost 25% or less of fecundity with exposure. Notably, ECA705 (#5) made 13% more offspring under exposure.

**Figure 2:**
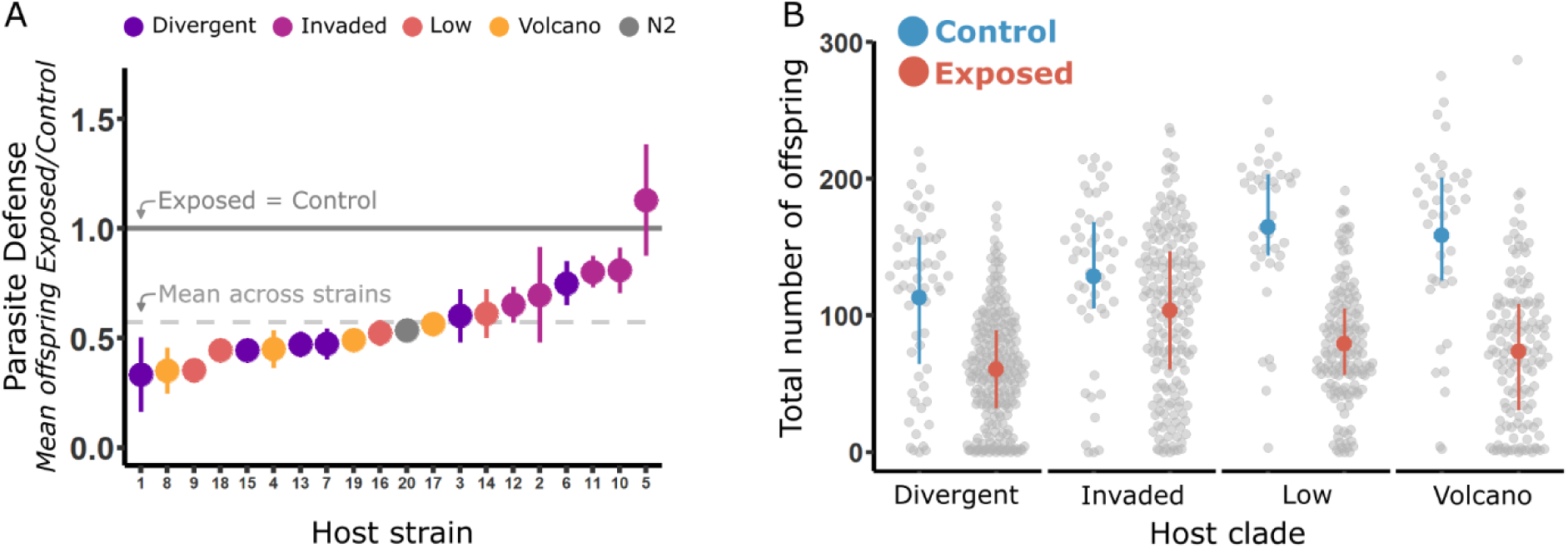
Host strains and groups vary in defense against parasites. **A** shows each host strain’s level of defense, with defense defined as the mean number of total offspring per host following exposure, divided by the mean number per host in the control treatment. Ratios below 1 (solid gray line) indicate that hosts made fewer offspring with parasite exposure relative to control. Error bars show the standard error of this ratio. The dotted gray line represents the average defense across host strains. Host strains are arrayed along the x-axis in order of increasing defense, and they are colored according to group, with the host strain N2 in gray. Host strains are indicated by their numeric ID (Table A in S1 Text, Fig. 1). **B** displays the fecundity of hosts in control (blue) and exposed conditions (red) by host group. Gray points show the total number of offspring for individual hermaphrodites. Filled points indicate the mean per treatment, and error bars show the interquartile ranges of the data. Exposed includes data from all four exposure treatments. Each host group is represented by 4-6 host strains. N2 is excluded from this plot.

Defense also varied with host phylogenetic group (Fig. 2a,b; Table E in S1 Text - interaction of host group and exposure: *χ^2^*=41.6, df=3, p<0.001). Exposure did not significantly reduce mean fecundity for hosts in the Invaded group (Tukey test: t = 1.84, df = 33, p = 0.601): they made 103.6 ± 4.3 offspring per host under exposure, relative to 128.5 ± 8.5 in the control treatment, a loss of only 19%. In contrast, exposure reduced fecundity by 54% in the Volcano group (158.5 ± 10.8 in control to 73.5 ± 4.5 in exposed). Fecundity losses were also substantial for the Divergent (-46%) and Low groups (-52%) (Tukey tests: p ≤ 0.003 in all three cases).

### Fitness effects of specific exposed treatments

The four exposure treatments varied in their impact on host fecundity (Fig. 3a, Table F in S1 Text). They each substantially reduced fecundity relative to the control treatment (Tukey tests: p < 0.001 in all four cases). Exposure to a low dose of *N. parisii* had the smallest effect, reducing fecundity by only 28% on average (107.1 ± 4.4 offspring per host, relative to 148.4 ± 4.9 per host in the control). Exposure to a high dose of *N. ironsii* had the largest effect, reducing fecundity by 57% on average, to 63.6 ± 3.1 offspring per host. Across parasite species, the effect of exposure increased with dose: hosts made an average of 95.9 ± 2.9 offspring at low doses and 73.5 ± 2.4 at high doses. Across doses, *N. ironsii* had a greater effect on fecundity than *N. parisii*: hosts made an average of 95.1 ± 2.9 offspring under exposure to *N. parisii* and 74.1 ± 2.4 under exposure to *N. ironsii*.

**Figure 3:**
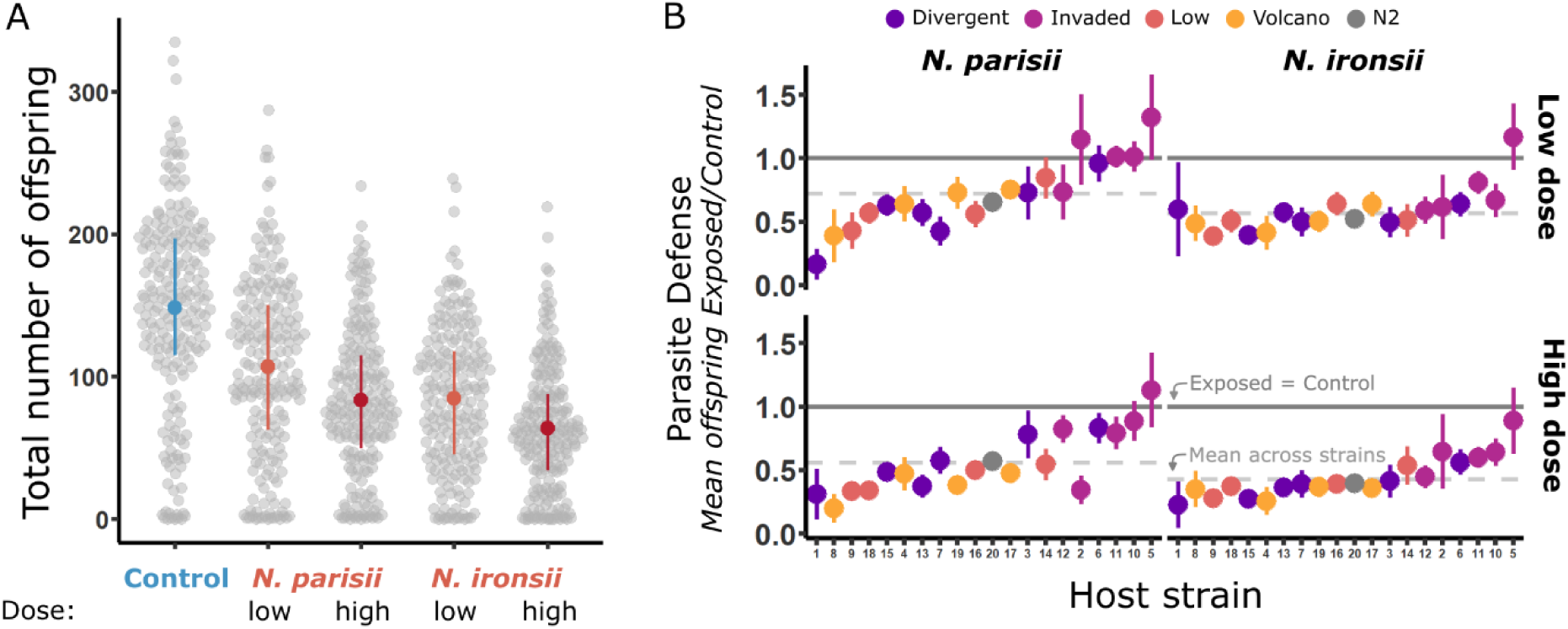
Parasite species and doses vary in their mean fitness effect. **A** displays the fecundity of hosts in control (blue) and exposed treatments (red). In exposed treatments, hosts encountered either *N. parisii* or *N. ironsii* at low (lighter red) or high (darker red) doses. Gray points show the total number of offspring for individual hermaphrodites. Filled points indicate the mean per treatment, and error bars show the interquartile ranges of the data. Each treatment is represented by 203-209 hosts across 20 strains. **B** shows each host strain’s level of defense for each exposure treatment, with defense defined as the mean number of total offspring per host following a specific exposure treatment, divided by the mean number per host in the control treatment. Ratios below 1 indicate that hosts on average made fewer offspring with parasite exposure relative to control. Error bars show the standard error of this ratio. The dotted gray line represents the average defense across host strains in that treatment. Host strains are arrayed along the x-axis in order of increasing overall defense, as shown in Fig. 2a, and they are colored according to group, with the host strain N2 in gray. Host strains are indicated by their numeric ID (Table A in S1 Text, Fig. 1).

For each exposure treatment, host strains showed continuous variation in defense, as shown by significant variation among host strains in fecundity loss from control under each exposure treatment (Fig. 3b, S2 Fig., Table G in S1 Text - significant interactions of host strain and treatment for all four comparisons). Estimates of broad sense heritability for fecundity in each exposed treatment ranged from 0.32 [0.15,0.51] under exposure to a high dose of *N. ironsii* to 0.47 [0.29,0.67] under exposure to a high dose of *N. parisii* (S3 Fig.).

We did not, however, see strong evidence for specific responses to dose or parasite species: across exposure treatments, changes in fecundity could be predicted by the additive effects of host strain and exposure treatment (Table H in S1 Text - insignificant interaction of host strain and exposed treatment: *χ^2^*=69.9, df=57, p=0.123). Consistent with this finding, host strains that were relatively defended in one exposure treatment tended to be relatively defended in other exposure treatments: defense estimates were significantly positively correlated between treatments in five of six comparisons (Fig. 3b, Spearman’s rank correlation: 0.51 ≤ *ρ* ≤ 0.80, Bonferroni-corrected α value = 0.008, Table I in S1 Text).

### Mechanisms of defense against parasites: life history

Variation in defense against parasites may arise from differences in host life history. Specifically, we hypothesized that strains that produce offspring earlier are more defended against fecundity loss from infection. To test this hypothesis, we examined variation in the timing of reproduction in the control treatment and related it to our estimates of defense.

The average control host produced most of its offspring on the second and third days of reproduction (50.6% and 34.2%, respectively) (Figure 4a: blue). Host strains varied in the timing of reproduction (S4 Fig., blue; Table J in S1 Text - interaction of host strain and day: *χ^2^*=425.1, df=76, p<0.001). For example, host strain QX1792 (#11) made 84.0% of its offspring in the first two days of reproduction, while QX1791 (#18) made only 36.4% of its offspring during the same period. Broad-sense heritability estimates for the number of offspring on individual days were comparable to the estimate for total number of offspring in the control treatment (H^2^ = 0.38 [0.20,0.59] for day 2, H^2^ = 0.33 [0.15,0.54] for day 3).

**Figure 4:**
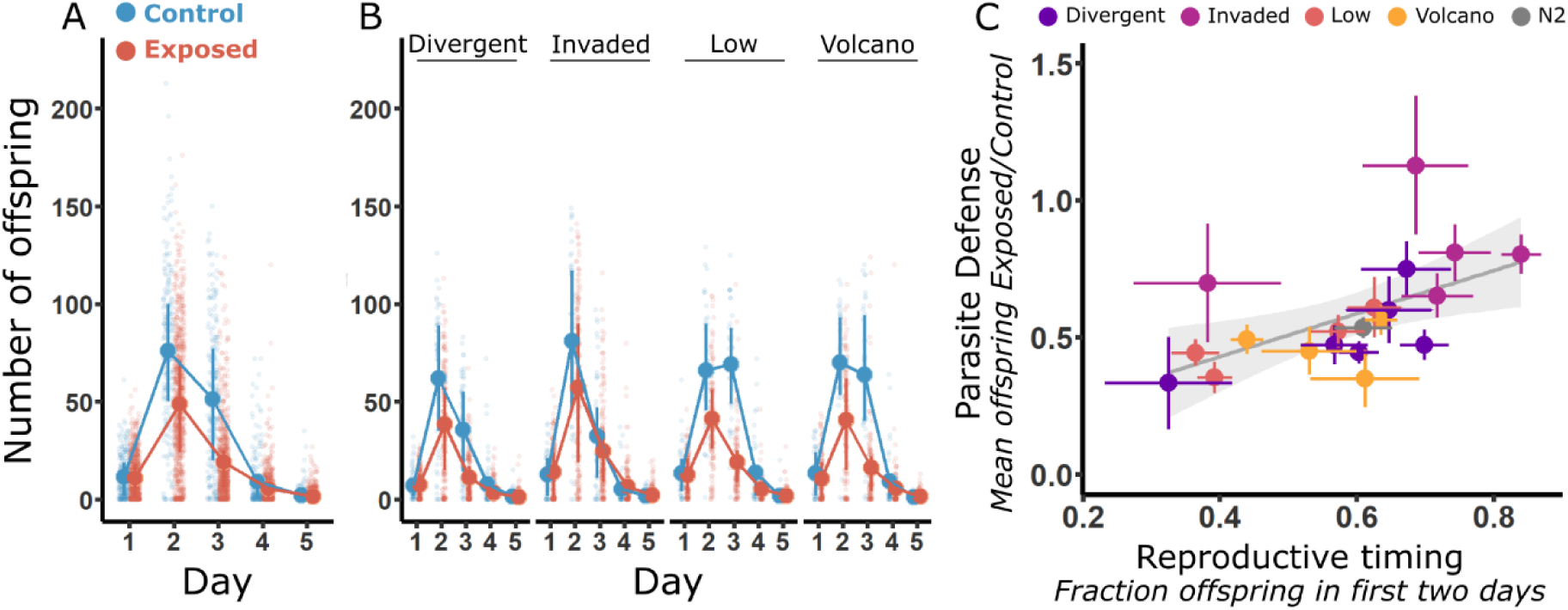
Host strains that reproduce early are more defended. **A** shows the number of offspring per host per day, for each of the five days of the fecundity assay across all host strains. Blue denotes hosts in the control treatment, and red denotes hosts in exposed conditions, representing all four treatments. Solid points show the mean, and shaded points show raw data from individual hermaphrodites. Error bars show the interquartile ranges of the data. Days capture reproduction during the prior 24 hours (day 1 = 0-24 hours, day 2 = 24-48 hours, day 3 = 48-72 hours, day 4=72-96 hours), with the exception of day 5 which captures reproduction from 96-144 hours. **B** shows these same data broken down by host group, with 4-6 host strains per group. **C** shows the relationship between reproductive timing and defense. Each point gives the mean value for a host strain, with standard errors, colored by host group. Reproductive timing is represented by the fraction of a host’s total offspring made on the first two days of reproduction in the control treatment; higher values indicate faster reproduction, because a greater fraction of offspring were made early. Defense against parasites is represented as in Fig. 2a.

Reproductive timing also varied by host group (Fig. 4b: blue; Table K in S1 Text - interaction of host group and day: *χ^2^*=103.1, df=12, p<0.001). For example, in the control treatment, hosts in the Invaded group reproduced relatively quickly, making 68.6% of their offspring in the first two days of reproduction. In contrast, hosts in the Low group only made 48.7% of their offspring in the first two days, with equivalent mean numbers of offspring on the second and third days of reproduction (day 2 = 66.1 ± 5.0 offspring per host; day 3 = 69.2 ± 4.6).

Exposure shifted the timing of reproduction (Fig. 4a: red; Table L in S1 Text – interaction of day and condition: *χ^2^*=444.0, df=4, p<0.001). On average across exposure treatments, parasite exposure reduced offspring number by 62.7% on the third day of reproduction. This impact was far larger than the 35.8% reduction observed on day 2, the other major day of reproduction. As a result, compared to the average control host, the average exposed host made relatively more of its offspring on the second day of reproduction (56.6%) and fewer on the third day (22.2%), when the fitness impact of parasite exposure was greatest (Fig. 4a,b).

Consistent with our hypothesis, host strains that reproduced earlier were more defended against parasites. Across strains, we found a positive relationship between defense and the fraction of offspring made on the first two days of reproduction in the control treatment (Fig. 4c, Table M in S1 Text: *β* = 0.78 ± 0.27, t value = 2.92, p=0.009). We found no relationship between defense and total fecundity (Table N in S1 Text: *β* = -0.0002 ± 0.0010, t value = -0.21, p=0.839).

### Mechanisms of defense against parasites: resistance

Variation in defense may also arise from differences in resistance, defined as a host’s ability to limit establishment and proliferation of an infection. Specifically, we hypothesized that strains that are more resistant are more defended against fecundity loss from infection. To test this hypothesis, we used fluorescence *in situ* hybridization (FISH) to estimate parasite prevalence, measured as the fraction of hosts that were infected following 48 and 72 hours of exposure to a low dose of *N. parisii*. We also estimated the infection load of individual infected hosts, which we measured as the fraction of host body area occupied by parasites. At 48 hours, we assayed 2,986 hermaphrodite hosts across 20 strains, with each strain independently tested in at least three replicate blocks with 49.0 ± 2.14 hosts per block. At 72 hours, we assayed 1,187 hermaphrodite hosts across 10 strains, each tested in 2-4 blocks with 34.9 ± 0.15 hosts per block (Table O in S1 Text). Host strains with lower infection prevalence and load were considered more resistant. We then related these estimates of resistance to our estimates of defense.

After 48 hours of exposure, an average of 81.1 ± 1.6% of hosts per replicate were infected. Host strains varied in their prevalence of infection (S5a Fig., Table P in S1 Text – effect of host strain on zero-inflation: *χ^2^*=84.5, df=19, p<0.001), ranging from a low of 67% of hosts infected for ECA1977 (#16) to a high of 96% of hosts infected for ECA744 (#19). Prevalence also varied among host groups (Table Q in S1 Text – effect of host group on zero-inflation: *χ^2^*=24.5, df=3, p<0.001): hosts from the Volcano group had slightly higher prevalence, 86.9 ± 2.5 %, than the other groups (Low = 74.8 ± 4.3%, Divergent = 79.2 ± 2.9%, Invaded = 81.9 ± 3.4%).

Infection load was low at 48 hours. The average load of infected hosts per replicate ranged from 0.17% to 9.0% of host body area fluorescent. Host strains varied in infection load at this early time point (S6 Fig., Table P in S1 Text – effect of host strain on conditional model: *χ^2^*=51.2, df=19, p<0.001). Infected hosts of CB4856 (#9) and ECA743 (#12) had relatively small loads (mean <1.4%), while infections of N2 (#20) and ECA744 (#19) were ∼ 3-fold larger (mean >4%). Infection load did not vary among host groups (Table Q in S1 Text – no effect of host group on conditional model: *χ^2^*=1.8, df=3, p=0.614).

After 72 hours of exposure, 99.6 ± 0.3% of hosts were infected (S5b Fig.). The average load of infected hosts per replicate was higher than at 48 hours, ranging from 3.7% to 25.2% of host body area infected. Host strains also varied in infection load at this later time point (Fig. 5a, Table R in S1 Text – effect of host strain: *χ^2^*=30.3, df=9, p<0.001). Hosts from strain ECA2334 (#13) had the smallest load – 7.4 ± 2.4% – while ECA1997 (#8) and CB4856 (#9) had the largest loads, at 15.2 ± 7.3% and 20.2 ± 3.1%, respectively. Infection load again did not vary among host groups (Table T in S1 Text – no effect of host group: *χ^2^*=4.5, df=3, p=0.216).

**Figure 5:**
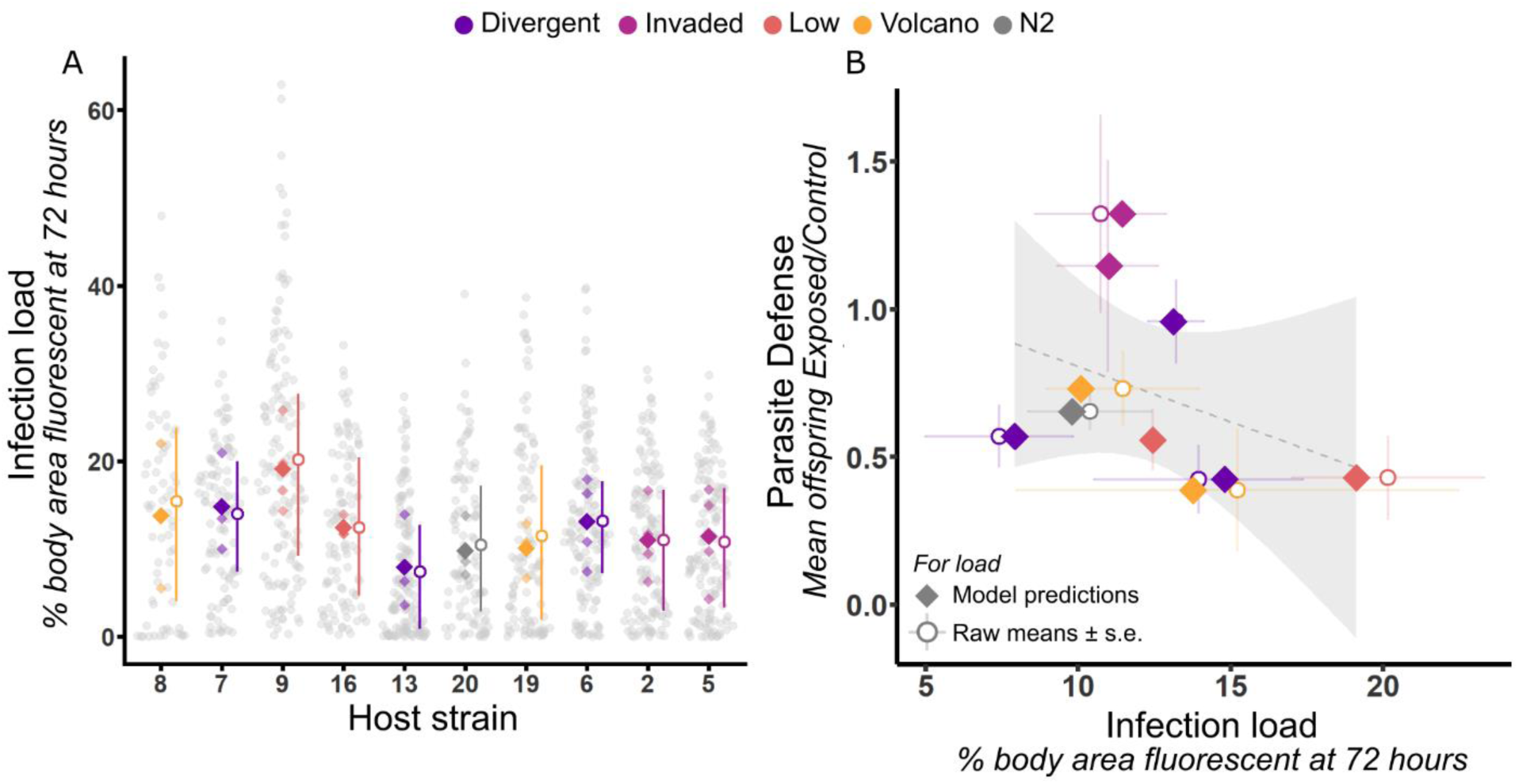
Defense against parasites does not correlate with measures of resistance. **A** presents infection load at 72 hours, measured as the percent of body area fluorescent in 2D images of hosts. Higher infection load is interpreted as lower resistance. Gray dots show infection load of individual hosts. Small shaded diamonds show model predicted strain means for replicates from the statistical model in Table R in S1 Text, and larger diamonds show the means of these model predictions. The open circles show means of the raw data plus interquartile ranges. Each host strain is represented by 71-143 hosts assayed across 2-4 blocks. Host strains are arrayed along the x-axis in order of increasing defense against a low dose of *N. parisii*, as shown in Fig. 3b, top left, and they are colored according to group, with the host strain N2 in gray. Host strains are indicated by their numeric ID. **B** shows the relationship between infection load at 72 hours and defense. Increasing values on the x-axis indicate decreasing resistance. Each point represents a host strain, colored by host group. Defense estimates reflect the response to a low dose of *N. parisii*, as in Fig. 3b, top left. For load, diamonds show the means of model predictions, as in A. Circles show the means of the raw data, with standard error across replicates.

Though we detected variation in infection load among host strains at both timepoints, we also observed substantial variation among individuals of the same strain (S6 Fig., Fig. 5a). Consistent with this individual variation, our estimates of broad-sense heritability for infection load were low: 0.022 [0.002, 0.046] at 48 hours, and 0.041 [0.000,0.097] at 72 hours.

Contrary to our hypothesis, host strains with lower loads (i.e., more resistant) were not more defended against parasites. Across strains, we found no relationship between defense and infection load at 48 (S6b Fig., Table Ua in S1 Text: *β* = -0.02 ± 0.13, t value = -0.16, p=0.878) or 72 hours (Fig. 5b, Table Ub in S1 Text: *β* = -0.04 ± 0.03, t value = -1.10, p=0.302).

## DISCUSSION

In order to understand how host populations might respond to parasite selection, we characterized genetic variation in defense against parasites and identified traits that covaried with defense. We found that Hawaiian strains of *C. elegans* varied substantially in defense against *Nematocida* parasites, ranging from as much as a 60% reduction in fecundity to no change in fecundity under parasite exposure. Strains from the Invaded group were notably more defended than strains from the other three Hawaiian groups. We did not find support for the hypothesis that heritable variation in resistance covaried with defense against parasites. Rather, heritable variation in host life history covaried with defense. Our results support the hypothesis that reproducing early protects hosts against fitness losses from parasites.

### Characterization of genetic variation in defense against parasites

Hawaiian isolates of *C. elegans* show extensive genetic diversity that is hypothesized to reflect patterns of ancestral genetic variation, prior to global spread of the species [11,47]. We accordingly expected to find substantial variation in defense among Hawaiian strains. This prediction was supported (Fig. 1b, <u>2a</u>). Though prior studies have not specifically investigated Hawaiian strains, our findings are consistent with prior work showing variation among wild *C. elegans* strains in their fitness in the presence of *Nematocida* [33,38,39]. Notably, in [40] the presence of *Nematocida* altered the outcome of competition among 22 different *C. elegans* strains, with certain strains increasing or decreasing in abundance more than expected based on control competitions. Broad surveys like our study and [40] capture among-population variation in *Nematocida* defense, leaving open the question of how much variation in defense exists within populations of *C. elegans*. The results of [33] argue strongly that there is also within-population variation in defense: the presence of *N. ausubeli* reversed the outcome of pairwise competitions among strains sampled from a single orchard. Collectively, these works support significant potential for the evolution of defense in response to selection by *Nematocida*.

Variation in defense could be maintained by costs or specificity of defense, among other things [49–51]. We found no evidence of costs – total fecundity and defense were not negatively correlated across host strains. Costs are, however, difficult to reject. Notably, in many host-parasite systems [52,53], including *C. elegans-Nematocida* [54], defenses are often costly only in specific contexts. We also found little support for specificity as a mechanism maintaining variation in defense. Strains shifted somewhat in their relative performance against *N. parisii* versus *N. ironsii* (Fig. 3b). This finding is consistent with [40], where individual host strains differed in their relative fitness against different *Nematocida* species, as measured by relative changes in abundance in multi-strain competitions. However, the overwhelming pattern in our data was a strong correlation in performance across exposure conditions, such that strains with relatively high defense against *N. parisii* also had relatively high defense against *N. ironsii* (Fig. 3b). We note that our approach does not directly test the genotype-by-genotype specificity that is hypothesized to maintain variation in antagonistic interactions [50,55,56]. To do so requires evaluating specificity at the intraspecific level, using different strains of the same parasite species. This test should be the subject of future work. Beyond costs and specificity, spatiotemporal variation in parasite selection is a viable explanation for the maintenance of variation in defense [57,58]. *C. elegans* disperses between ephemeral patches of rotting vegetation, suggesting that selection by abiotic and biotic forces is highly heterogeneous in time and space [59]. Data on the incidence and distribution of *Nematocida* across habitat patches would help evaluate this hypothesis.

We did not *a priori* hypothesize that the four Hawaiian phylogenetic clusters would show distinct patterns of defense against parasites, so the finding of substantially elevated defense in the Invaded group was a surprise to us (Fig. 2b). Strains classified as members of the Invaded group bear signatures of admixture with non-Hawaiian lineages, and small portions of their chromosomes contain “swept” haplotypes common in non-Hawaiian strains [47]. These haplotypes reflect recent selective sweeps that are thought to be associated with the global spread of *C. elegans* [11,60]. Strains with substantial representation of swept haplotypes have elevated fecundity and earlier reproduction [46]. Our data suggest that these swept haplotypes may also confer increased defense against parasites. Targeted comparison of swept and non-swept strains is needed to evaluate this hypothesis.

### Tests of traits that covary with defense against parasites

Given this genetic variation in defense among hosts, we then asked: which heritable traits make some hosts more defended than others? Because we broadly compared the relative fitness of host strains under parasite exposure, irrespective of the infection status of individual hosts, many factors could have contributed to variation in defense among host strains, including behavior, overall vigor, and within-host processes. We specifically tested two hypotheses, that 1) life history traits and/or 2) resistance traits covaried with defense. In support of the life history hypothesis, we found that the most defended host strains were those that reproduced quickly, such that relatively more eggs were laid on the first two days of reproduction (Fig. 4c). This result is consistent with life history theory predicting that reproducing early should be advantageous in the face of forces, like parasites, that reduce opportunity for future reproductive success [19,20]. We do not see clear evidence that parasite exposure induced accelerated reproduction in *C. elegans*, as has been observed in snails [22], crickets [23], *Daphnia magna* [61,62], and house sparrows [63]. Exposed hosts produced a greater *fraction* of their offspring in the first two days, but this shift seemed to reflect the strong negative effect of parasites on offspring number in later days (notably on day 3), rather than an increase in the number of offspring produced early. Indeed, exposed hosts did not produce a greater *number* of offspring than control hosts on the first two days of reproduction (Fig. 4ab, S4 Fig.). Thus, defense against parasites is associated with constitutively early reproduction, a pattern that has also been observed in perch [64] and snails [24,25,65].

It is possible that reproductive timing functions as a form of tolerance in *C. elegans* hosts. If so, we would expect the fitness of early-reproducing host strains to be less sensitive to increasing parasite load than that of late-reproducing host strains. Alternately, reproductive timing may be correlated with other traits that confer defense, such as host vigor; more defended host strains may simply be those in better condition and thus able to reproduce more quickly [15,66]. Better diets do accelerate reproduction in *C. elegans* [54,67], supporting the link between vigor and reproductive timing. However, those strains that produced the greatest total number of offspring (e.g., N2 - #20, ECA744 - #19) were not the most defended (Fig. 1b, 2b, Fig. S4). Directly testing the tolerance hypothesis, and ruling out vigor, requires estimates of host fitness at several parasite doses to measure the rate of change in fitness with increasing parasite load [15]. This test would ideally isolate the effect of variation in reproductive timing by comparing host strains with distinct life histories but similar genetic backgrounds [e.g., 68].

Our results suggest that reproductive timing could respond readily to parasite selection, as we found a strong heritable component of variation in daily fecundity, consistent with prior studies [46,69]. By this argument, parasite selection could specifically favor host strains with swept haplotypes, which are associated with earlier reproduction [46]. However, reproductive timing is a fundamental life history trait that we expect to be under strong selection by many factors, including extrinsic mortality and resource availability [70–74]. Given this, early reproduction should not be viewed as a specific adaptation to *Nematocida*, and we think it unlikely that the variation in reproductive timing observed in our wild host strains can be explained by historical patterns of parasite selection. Rather, our results indicate that the observed variation in defense among our wild strains is associated with variation in life history that likely arose due to other evolutionary forces acting in natural populations. Moreover, we expect it may be challenging to detect cases of direct parasite-mediated selection on reproductive timing in natural populations of *C. elegans*, given the diverse selective agents, genetic constraints, and remarkable plasticity of life history traits in this species [rev. in 75]. Parasite-mediated selection on life history traits may only be detectable in experimental contexts, where confounding factors can be controlled.

We expect life history traits to nonetheless have major implications for the evolution of defense in wild populations of *C. elegans*. Hosts in the plant genus *Silene* provide an illustrative example: annual species rarely become infected with the sterilizing fungus *Microbotryum,* while perennial species are infected at high rates [28]. Yet under direct inoculation, annual species have lower resistance to infection than perennial species. This pattern suggests that evolutionary shifts in *Silene* life history – from the perennial to annual habit – indirectly provided defense against infection and thereby selected against the maintenance of costly resistance [29]. Likewise, factors that promote earlier reproduction in wild populations of *C. elegans* could reduce the strength of parasite-mediated selection, thereby relaxing selection on other traits directly associated with *Nematocida* defense. Thus, our findings add to a large body of work supporting host life history as an organizing principle in host-parasite interactions and defense evolution [76–79].

Our results do not support the alternative hypothesis that resistance traits explain variation in host defense. The most defended host strains did not have relatively low infection prevalence or load, at either 48 or 72 hours post-exposure, and strains with relatively low burden had similar defense levels to strains with relatively high burden (Fig. 5b, S6b Fig.). This result echoes many prior studies: the relative fitness of *Arabidopsis thaliana* accessions was uncorrelated with growth of *Pseudomonas syringae* following exposure [14]. Similarly, *P. aeruginosa* load did not correlate with variation in survival post-infection of lines of *Drosophila melanogaster* [80], and increases in load of *Listeria monocytogenes* did not explain increased parasite-mediated mortality of *D. melanogaster* mutants [81]. Our findings do not reject the possibility for resistance to promote host fitness in the face of parasites – indeed, prior studies of *Nematocida* and *C. elegans* find that strains with reduced parasite establishment and proliferation can have elevated fitness [39,40,43]. Rather, our findings argue that resistance traits are not the dominant explanation for the variation in defense observed among our panel of wild host strains. We note, however, that we had limited power to reject an association between resistance traits and defense, because of the lower number of host strains at 72 hours post-exposure, when infection load was most variable. It would be worth evaluating more strains at this and later time points of infection. Nevertheless, our primary resistance trait – parasite load – showed far more variation within host strains than between strains, resulting in a very low estimate of heritability [see also 82,83]. We expect load to be even more variable outside the controlled conditions of our artificial exposure. Thus, even if load did covary strongly with defense, we predict limited potential for this resistance trait to respond to parasite selection.

### Future directions

Our analyses compared single-generation measures of fecundity and infection to make inferences about the average defense level of hosts of different genetic backgrounds. *C. elegans* is a short-lived invertebrate that reproduces largely by selfing on ephemeral bacterial blooms. Persistence of a selfing lineage ultimately requires the export of dispersal larvae that can successfully colonize new patches in a metapopulation [33,59,84]. Within a patch, *Nematocida* transmits readily from parent to offspring via spores shed in the environment [36], and infection prevalence and load can escalate quickly as a host lineage proliferates [85]. Thus, transmission between generations may be an important consideration in parasite selection and defense evolution in this system. Indeed, there is evidence for transgenerational immunity against *Nematocida*, such that offspring of exposed parents can be much more resistant than offspring of healthy parents [83]. Our single-generation approach does not account for these multigenerational effects. Genetic variation for defense is very likely sensitive to the scale at which defense is measured [38]. For example, in our single-generation assays, the host strain CB4856 (ID #9) ranked as poorly defended relative to N2 (ID #20) (Fig. 2a). However, over the course of multi-generational epidemics, CB4856 maintained higher fitness and lower parasite loads than N2 [85], consistent with prior studies reporting CB4856 as relatively resistant and defended [39,40]. These differences raise the important hypothesis that the most successful defense strategy depends on the level of selection [86,87], and this tension could contribute to maintaining the variation in defense we observed in this study [88–91].

## METHODS

We first conducted a fitness assay that measured the daily fecundity of hosts under control and exposed conditions. These data were used to characterize defense against parasites and life history of 20 host strains. We conducted a separate assay to measure resistance traits of these host strains. In all experiments, experimental conditions were fully blinded during data collection.

### Host strains

The 19 Hawaiian *C. elegans* strains in this study each represented a genetically distinct isotype, defined by [47] as a collection of strains sharing >99.9% of single nucleotide variants. Each strain was established by a single hermaphrodite. To ensure wide representation of genetic variation across the species, we used the relatedness clusters reported in [47] - Divergent, Invaded, Low, and Volcano - to guide selection of these strains. Each of the four groups was represented by 4-6 strains collected across three Hawaiian islands, with the exception of the Volcano group which is only found on the island of Hawai’i (S1 Fig.). With the well-characterized lab strain N2, our total number of strains was 20 (Table A in S1 Text).

We obtained all strains from CaeNDR, including N2, and archived them at −80°C following CaeNDR guidelines [92]. Prior to an experimental block, we thawed frozen host strains and reared them for two weeks (∼4 generations) at 20°C on Nematode Growth Medium (NGM Lite, US Biological) plates seeded with *Escherichia coli* strain OP50. We added 20% agarose to the media to reduce burrowing beneath the surface of the plate (NGMA). All experiments were similarly conducted at 20°C on OP50-seeded NGMA plates. OP50 was obtained from the *Caenorhabditis* Genetics Center and cultured in LB broth at 37°C for 48 hours prior to use.

### Parasite species

We used two species of *Nematocida* to determine if host defenses are general or specific to a given parasite species. The two species, *N. parisii* and *N. ironsii,* are closely related but appear to be distinct ecologically: *Nematocida parisii* is commonly found infecting *C. elegans* and *C. briggsae* and has largely been sampled in Europe [31], while *N. ironsii* was isolated once from *C. briggsae* in Kauai, Hawaii [93]. For *N. parisii*, we used strain ERTm1, the original strain isolated from *C. elegans* in France [36]. For *N. ironsii,* we used strain ERTm5, the only available strain. Both species are transmitted horizontally; there is no vertical transmission [36].

For each species, we generated spore stocks for experimental exposures and quantified their spore concentration as described in [38]. We also generated parasite-free lysates for use in the control treatment, as in [38]. To identify comparably infective doses of *N. parisii* and *N. ironsii* stocks, we exposed 1,200 hosts of the strain N2 to a range of doses of each stock and measured their infectivity after 3 hours. Specifically, we measured the average number of infective spores encountered by a host using FISH to label viable spores within the host intestine (see detailed methods in section *Measuring Parasite Resistance*) [36,39]. From these measurements, we estimated the volume of each spore prep for which we expected an average of 1.5 and 3 infective spores encountered per host after 3 hours of exposure. The prior estimate was used as the volume for low dose exposures and the latter as the volume for high dose exposures in the fitness assay described below. We found our *N. ironsii* stock to be more infective than our *N. parisii* stock, so hosts were exposed to fewer *N. ironsii* spores at each dose (Table B in S1 Text).

### Fitness assay

Daily fecundity was measured on individual hermaphrodites subjected to five different treatments, one control and four exposed. For the four exposure treatments, hermaphrodites were exposed to a low or high dose of one of the two parasite species (Table B in S1 Text). For the control treatment, parasite inoculum was replaced with a volume of control lysate equivalent to our highest exposure volume (63.8 µL) to conservatively account for any effects of the spore solution beyond the parasites themselves [e.g., 94]. In exposure treatments, our fecundity estimates were purposefully blind to the infection status of individual hosts. We accordingly measured the expected relative fecundity of a host of a given genetic background (i.e., strain) reared in the presence of parasites. Many factors can potentially contribute to defense against parasites (e.g., avoidance, resistance, tolerance), so by generally estimating fecundity in the presence of parasites, irrespective of infection status, we were able to estimate the net fitness effect of these factors, without biasing our estimates towards one particular strategy or mechanism [14].

We modeled our fitness assay on [46]. To begin the assay, we thawed a frozen vial of a given host strain, allowed hosts to proliferate for four generations at 20°C, then collected eggs using a standard bleach wash. Eggs were plated directly on *E. coli* to prevent starvation. Twenty-four hours later, we collected first-stage (L1) larvae in M9 buffer and measured their concentration. To establish each treatment, we suspended 200 L1 hosts with a mixture of 150 µL of concentrated OP50, the treatment-specific volume of spores or control lysate (Table B in S1 Text), and M9 buffer to reach a volume of 500 µL. This suspension was spread evenly across a 60mm plate of NGMA and allowed to dry before storing at 20°C for 24 hours. After 24 hours of exposure, hosts had reached the L4 larval stage, just before adulthood and egg-laying. We collected L4 hosts and washed them to remove external spores. Hermaphrodites were picked individually to 35mm NGMA plates seeded with 25 µL of OP50. We allowed hermaphrodites to reach reproductive maturity and lay eggs on this first plate for 24 hours. We then moved each hermaphrodite parent to a new 35mm plate to continue laying eggs. We repeated this transfer every 24 hours for a total of 96 hours (4 days). After 96 hours, few eggs were being laid, so we moved each original parent once more and allowed them to lay eggs for an additional 48 hours (144 hours total). We recorded whether the parent was alive or dead during transfer and ceased transferring if it was identified as dead. We censored males (n=36, 3.3%) and hermaphrodites that disappeared, burrowed into the agar, or were lost (n=36,3.3%). Table C in S1 Text further details sample size and block structure.

To count viable offspring, 35mm plates of eggs were incubated at 20°C for 48 hours after removal of the parent. During this time, eggs hatched and offspring grew large enough to be imaged. We imaged entire plates using a custom imaging platform modeled on [95] and [46]. We used a DMK33GP031 camera (Fujinon HF255A-1 25 mm lens) and IC Capture software from the Imaging Source (Charlotte, NC, USA). For each hermaphrodite, we imaged plates of offspring produced at 24 (day 1), 48 (day 2), 72 (day 3), 96 (day 4) and 144 hours (day 5+) of reproduction (> 5,000 plates total). Each plate was imaged twice to facilitate identification of offspring. To quickly and consistently count the number of offspring on each plate, we used a custom computer vision pipeline trained to recognize juvenile hosts [96]. For the final estimate for each plate, we used the average of the computer vision counts for the two separate images of the plate. Computer vision results were extensively checked against manual counts, which were done using ImageJ’s Multi-point Tool [97]. Computer vision counts were highly correlated with manual counts but substantially more consistent. Manual counts notably varied significantly with the identity of the counter [96].

### Resistance assays

We characterized variation in resistance by estimating each strain’s infection prevalence and load after exposure. We focused on resistance to a low dose of *N. parisii*, as we saw substantial variation in defense among host strains under this treatment and were able to maximize the number of host strains tested with the same spore stock. Block structure and sample size are detailed in Table O in S1 Text.

To facilitate comparison with fitness data, hosts were raised in a manner similar to the fitness assay. Twenty-four hours after extracting eggs, we suspended 2,000 L1 hosts of a given strain with a mixture of 400 µL of concentrated OP50, 14.7 µL of *N. parisii* spore stock, and M9 buffer to reach a volume of 1 mL. This suspension was spread evenly across a 60mm plate of NGMA and allowed to dry before storing at 20°C for either 48 or 72 hours. These time points were chosen to span the period during which the infection establishes and proliferates. At 48 or 72 hours, hosts were collected in M9, washed twice to remove spores and *E. coli*, then fixed in acetone. We used FISH to quantify the size of host’s infection. Specifically, we used a fluorescent probe to stain exposed hosts for a *Nematocida* ribosomal RNA small subunit sequence [36]. Hosts were mounted and imaged at 10x magnification with a fluorescent microscope (DM6 B, Leica) and Texas Red filter. Using ImageJ [97], we cast each image in black and white, outlined a host’s body to measure its area, then set the threshold brightness to highlight the fluorescent areas of the body (i.e., the areas with parasites). We used the function “Summarize” to calculate the percentage of the host’s body that was fluorescent. Male hosts were not included in analyses to facilitate comparison to the fecundity data, which was limited to hermaphrodites.

### Analyses

We ran analyses in R (v.4.4.0) using RStudio (v2024.04.0+735) [98]. We processed data and generated graphics using the package *tidyverse* [99]. We fit models using the packages *lme4* [100]*, glmmTMB* [101], and *survival* [102] (see S2 Text). We assessed model fits using the package *performance* [103]. We used likelihood ratio tests to compare models with and without terms of interest and Tukey’s tests in the package *emmeans* to compare specific factors levels [104].

### Fecundity and defense against parasites

We calculated a host’s total fecundity and fit linear mixed effect models. The data approximated a normal distribution because of the large range (0 to 335 total offspring per host), and we found a gaussian distribution provided a better fit than count distributions. We did not find evidence for variation across blocks in the fecundity of the strain N2, which was tested in every block (likelihood ratio test of the fixed effect of block on N2 fecundity: *χ^2^*=8.69, df=5, p=0.122). Nonetheless, we included block as a random effect, except in models examining variation among Hawaiian groups, because these models did not include N2. We also included a host’s plate of origin as a random effect in models comparing fecundity in control versus exposed conditions; in these analyses, exposed hosts came from four different exposure treatments, and this term accounted for the non-independence of hosts from the same exposure treatment/plate. We confirmed that our results were qualitatively unchanged when we excluded the strain XZ1514, which had very low fecundity.

To broadly evaluate variation in defense, we fit a model with host strain, parasite exposure (yes/no), and their interaction as fixed effects predicting the total fecundity of individual hosts. We compared this model to one without the interaction to determine if host strains varied in the effect of parasite exposure on fecundity. We repeated this analysis substituting host group for host strain and instead including host strain as a random effect.

We then evaluated the effects of specific exposure treatments on fecundity. To broadly compare treatments, we fit a model with host strain, treatment (n=5), and their interaction as fixed effects predicting total host fecundity. We then fit four separate models comparing fecundity of control hosts to hosts in each exposure treatment. We compared each model to a comparable model without the interaction of host strain and treatment to verify that the effect of parasite treatment varied across host strains. To evaluate the specificity of host responses to exposure, we fit a model to the total fecundity of individual hosts in exposure treatments only (i.e., control hosts excluded). Host strain, exposure treatment, and their interaction were included as fixed effects. We again compared this model to one without the interaction. Support for the interaction would indicate that the fecundity rankings of host strains changed substantially across exposure treatments, consistent with specificity. To further evaluate specificity, we quantified defense of each host strain as the ratio of the mean number of offspring in an exposure treatment to the mean number of offspring in the control treatment. Higher values indicated greater defense, because hosts retained more of their baseline fecundity under exposure. We then performed all possible pairwise correlations of defense scores across the four exposure treatments. A lack of positive correlation across treatments would suggest that host defense was specific, with for example a host relatively defended against one parasite species but not another. We used Spearman’s rank-order correlation for these tests, because defense estimates were not consistently normally distributed. We used Bonferroni correction to correct for multiple comparisons (n=6, alpha value = 0.008).

### Life history

Life history analyses included only hosts that reproduced (i.e., number of viable offspring > 0). Random effects were as described above, with the addition of individual host to account for repeated measures of fecundity through time. We first fit a linear mixed effect model to the number of offspring per host per day, from days 1-5, in the control treatment. We included host strain, day of reproduction, and their interaction as fixed effects and compared this model to one without the interaction to determine if host strains varied in when they made offspring. We repeated this analysis, substituting host group for strain as above. We then compared reproductive schedules under control and exposed conditions by fitting a model to daily fecundity counts with exposure (yes/no), day of reproduction, and their interaction as fixed effects, and host strain as an added random effect. Support for the interaction would indicate that control and exposed hosts varied in when offspring were made.

To test the hypothesis that early-reproducing host strains are more defended, we used linear regression to determine if there was a positive relationship across strains between defense and reproductive timing. Reproductive timing was calculated as a host strain’s mean fraction of total offspring per host made on the first two days of reproduction in the control treatment. Higher values indicated earlier reproduction. Defense against parasites was calculated as the ratio of a host strain’s mean total offspring per host under parasite exposure (all treatments) to the mean total per host in the control. We calculated both reproductive timing and defense from raw data, because we found no evidence of block effects for daily (random effect of block in Table J in S1 Text model: *χ^2^*=0, df=1, p=1.00) or total fecundity (random effect of block in Table D in S1 Text model: *χ^2^*=0, df=1, p=1.00). We asked whether our results could be explained by variation in overall fecundity by testing for a positive relationship across strains between defense and mean total number of offspring per control host.

### Resistance

We used infection prevalence and load at 48 and 72 hours post-exposure as proxies for resistance. Prevalence is the fraction of hosts infected after exposure; hosts were scored as 0 (no fluorescence = not infected) or 1. Load quantifies the extent of infection in infected hosts, and we measured it as the fraction of body area fluorescing.

For all models, we included host strain or group as a fixed effect, and experimental block and replicate within block as random effects. For models with host group, host strain was also included as a random effect. At 48 hours, a substantial number of hosts were not infected (0, no fluorescence). We accordingly evaluated variation among hosts in both prevalence and load by modeling a host’s fraction of body area fluorescing at 48 hours using a zero-inflated beta distribution. Structural zeros indicate individuals that are uninfected, while the conditional portion of the model represents the parasite load of infected individuals. At 72 hours, we excluded the few uninfected hosts (5/1187) and solely evaluated variation in load by modeling the fraction of body area fluorescing using a beta distribution. We compared models to comparable models without host strain/group to determine if hosts varied in resistance. For 48-hour models, we made this comparison for both the zero-inflated and conditional components.

To test the hypothesis that more resistant host strains are more defended, we used linear regression to determine if there was a negative relationship across strains between defense and infection load. Mean load for a host strain was the mean fraction of body area fluorescent, predicted from the statistical models above to account for significant block effects (random effect of block in 48h model, Table P in S1 Text: *χ^2^*=20.89, df=1, p<0.001; in 72h model, Table R in S1 Text: *χ^2^*=43.62, df=1, p<0.001). Predictions at 48 hours were derived from the conditional component of the model. Defense was calculated as above, substituting mean total offspring under exposure to a low dose of *N. parisii,* as this was the specific treatment applied for resistance measurements. Our results were qualitatively unchanged when we used raw estimates of mean load, as well as defense estimated across all exposure treatments.

### Heritability

Heritability of a trait defines its potential to respond to selection. We accordingly estimated broad-sense heritability of traits by quantifying the fraction of phenotypic variation attributable to variation among host strains. We used *MCMCglmm* [105] and *QGglmm* [106,107] because some of our traits had non-gaussian distributions. We included random effects as presented in the models above and used weakly informative priors. We assumed a gaussian distribution for fecundity traits. We assumed a Poisson distribution for percent of host body area fluorescent, rounding up to the nearest integer. We ran MCMC algorithms for 100,000 iterations, with a burn-in of 10,000.

## DATA AVAILABILITY STATEMENT

All data and analysis code associated with this study are publicly available at University of Virginia Dataverse at https://doi.org/10.18130/V3/U3KH8P.

## ACKNOWLEDGEMENTS

We thank members of the Gibson lab, Janis Antonovics, and Katja Kasimatis for helpful feedback on the project and the manuscript. Erik Andersen provided valuable guidance on the design, and Emily Troemel provided parasite strains. *Escherichia coli* was provided by the *Caenorhabditis* Genetics Center. Host strains were provided by the *Caenorhabditis* Natural Diversity Resource (CaeNDR; caendr.org).

## CAPTIONS FOR SUPPLEMENTAL FILES

**S1 Fig: Host strains included in this study. A** shows the sampling locations of the 19 Hawaiian strains. Points are jittered to reduce overlap. The 20^th^ strain, N2, is not from Hawaii and is not shown. The map was generated using the package *ggmap* [108]. The map tiles are by @ Stamen Design, @ Stadia Maps, and @ OpenMapTiles under [CC BY 4.0](https://creativecommons.org/licenses/by/4.0/). Data by [OpenStreetMap](https://www.openstreetmap.org/), under [ODbL](https://www.openstreetmap.org/copyright). **B** shows a neighbor-joining network of the 20 strains included in this study, plus four non-Hawaiian strains (in italics) that were not included in this study but are shown for added phylogenetic context. Colors of points in A and taxa in B correspond to the four relatedness clusters (i.e., group: Divergent, Volcano, Invaded, and Low). The neighbor-joining network was generated using [47]’s Figure 5 VCF dataset as the base variant call set. Three strains not included in that dataset (ECA1997, ECA1977, ECA2334) were downloaded from CaeNDR as VCF variant data and filtered to retain comparable sites. The additional strains were harmonized with [47]’s call set to generate a combined SNP matrix for the selected taxa. The merged VCF was converted to a NEXUS-formatted SNP alignment using vcf2phylip.py and then used to generate the neighbor-joining network in SplitsTree4 [109]. Table A in S1 Text provides further details on the host strains.

**S2 Fig: Fecundity of host strains across treatments.** The total number of offspring per host is shown for each of the five treatments, faceted by host strain. Points show raw data for total fecundity, while the lines show estimated marginal means from the full linear mixed effects model in Table F in S1 Text. For the raw data, unfilled points show the total number of offspring for individual hermaphrodites, and filled points show the mean. Error bars show 95% confidence intervals. For model estimates, solid lines indicate the mean and dashed lines the 95% confidence interval. Host strains are arrayed from top left to bottom right in order of increasing overall defense against *Nematocida*, as shown in Fig. 2a, and they are colored according to host group, with the host strain N2 in gray. Host strains are indicated by both their strain name and numeric ID.

**S3 Fig: Variation in fecundity within treatments.** Bars show the mean number of offspring per host, with 95% confidence intervals, for each host strain. Within a treatment, host strains are ordered from left to right by increasing fecundity in baseline control conditions. Host strains are indicated by their numeric ID (Table A in S1 Text, Fig. 1).

**S4 Fig: Reproductive schedules by host strain and exposure.** As in Fig. 4a,b, solid points indicate the mean number of offspring per host per day, and shaded points show raw data for individual hermaphrodites. Error bars show the interquartile ranges of the data. Blue denotes hosts in the control treatment, and red denotes hosts in exposed conditions, representing all four treatments. Host strains are arrayed from top left to bottom right in order of increasing overall defense against *Nematocida*, as shown in Fig. 2a. Host strains are indicated by both their strain name and numeric ID.

**S5 Fig: Infection prevalence by host strain** at (**A**) 48 and (**B**) 72 hours. Circles indicate the mean prevalence of infection across replicates, and error bars show interquartile ranges of the data. Host strains are arrayed along the x-axis in order of increasing defense against a low dose of *N. parisii*, as shown in Fig. 3b, top left, and they are colored according to host group, with the host strain N2 in gray. Host strains are indicated by their numeric ID.

**S6 Fig: Infection load at 48 hours and defense against parasites. A** presents infection load at 48 hours, measured as the percent of body area fluorescent in 2D images of hosts. Higher infection load is interpreted as lower resistance. Gray dots show infection load of individual hosts. Filled circles show means of the raw data plus interquartile ranges. Open circles show medians of the raw data. Host strains are arrayed along the x-axis in order of increasing defense against a low dose of *N. parisii*, as shown in Fig. 3b, top left, and they are colored according to host group, with the host strain N2 in gray. Host strains are indicated by their numeric ID. **B** shows the relationship between infection load at 48 hours and defense. Increasing values on the x-axis indicate decreasing resistance. Each point represents a host strain, colored by host group. Defense is given for the response to a low dose of *N. parisii*, as in Fig. 3b, top left. For load, diamonds show the means of model predictions for replicates from the conditional model in Table P in S1 Text.

**S1 Text:** Supplemental tables

**S2 Text:** Supplemental analysis of survival

## Notes

https://doi.org/10.18130/V3/U3KH8P

### Competing Interest Statement

The authors have declared no competing interest.

### Summary of Updates

Updated following reviewer comments. Portions of the Abstract, Introduction, and Discussion have been modified to incorporate alternative interpretations.

### Funding

- National Institutes of Health: R35GM137975

