## Supplemental Materials for "Defense against parasites covaries with reproductive timing, not with resistance"

### S1 TEXT – SUPPLEMENTAL TABLES


I. Table A: Host strains
II. Table B: Details of spore preps and doses for each parasite species
III. Table C: Number of hermaphrodite hosts per strain and treatment in the fitness assay.
IV. Table D: Statistical analysis of total fecundity in control and exposed conditions across host strains
V. Table E: Statistical analysis of total fecundity in control and exposed conditions across host group
VI. Table F: Statistical analysis of total fecundity across treatments
VII. Table G: Statistical analysis of change in total fecundity from control for each exposure treatment
VIII. Table H: Statistical analysis of total fecundity across exposed treatments
IX. Table I: Correlation of defense across exposure treatments
X. Table J: Statistical analysis of daily fecundity by host strain in the control treatment
XI. Table K: Statistical analysis of daily fecundity by host group in the control treatment
XII. Table L: Statistical analysis of daily fecundity in control and exposed conditions
XIII. Table M: Linear regression of reproductive timing and defense against parasites
XIV. Table N: Linear regression of fecundity and defense against parasites
XV. Table O: Replication structure for resistance assays
XVI. Table P: Statistical analysis of infection prevalence and load at 48 hours.
XVII. Table Q: Statistical analysis of infection prevalence and load by group at 48 hours.
XVIII. Table R: Statistical analysis of infection load at 72 hours.
XIX. Table T: Statistical analysis of infection load by group at 72 hours.
XX. Table U: Linear regression of resistance and defense against parasites

**Table A: Host strains**

| Strain/isotype | Group | Latitude | Longitude | Island | ID |
|---|---|---|---|---|---|
| CB4856 | Low | 21.330 | -157.860 | O'ahu | 9 |
| DL238 | Low | 19.110 | -155.810 | Hawai'i** | 14 |
| ECA1977 | Low* | 19.120 | -155.820 | Hawai'i | 16 |
| QX1791 | Low | 20.634 | -156.394 | Maui | 18 |
| ECA372 | Invaded | 22.122 | -159.664 | Kaua'i | 10 |
| ECA705 | Invaded | 20.039 | -155.439 | Hawai'i | 5 |
| ECA743 | Invaded | 20.742 | -156.323 | Maui | 12 |
| ECA812 | Invaded | 20.040 | -155.442 | Hawai'i | 2 |
| QX1792 | Invaded | 20.706 | -156.355 | Maui | 11 |
| ECA2334 | Divergent* | 22.133 | -159.651 | Kaua'i | 13 |
| ECA347 | Divergent | 19.439 | -155.304 | Hawai'i | 15 |
| ECA363 | Divergent | 20.724 | -156.304 | Maui | 7 |
| ECA724 | Divergent | 19.439 | -155.304 | Hawai'i | 6 |
| ECA740 | Divergent | 20.670 | -156.339 | Maui | 3 |
| XZ1514 | Divergent | 22.149 | -159.668 | Kaua'i | 1 |
| ECA1997 | Volcano* | 19.720 | -155.949 | Hawai'i | 8 |
| ECA730 | Volcano | 19.424 | -155.222 | Hawai'i | 4 |
| ECA744 | Volcano | 19.719 | -155.950 | Hawai'i | 19 |
| ECA746 | Volcano | 19.115 | -155.819 | Hawai'i | 17 |
| N2 | Non-Hawaiian | 51.450 | -2.590 | Great Britain | 20 |

* indicates group was assigned based on the species tree available on CaeNDR [20250625 release]. For all other strains, group was determined directly from assignments in [1], Figure 5, and verified in the latest species tree release. Geographic information was obtained from CaeNDR.
** Island of Hawai'i (Big Island)

**Table B: Details of spore preps and doses for each parasite species.**

| Parasite | Concentration | Low Dose | High Dose |
|---|---|---|---|
| *N. parisii* (ERTm1) | 61,143 spores/μL | 14.7μL, ~8.99 x 10^5 spores | 33.1μL, ~ 20.24 x 10^5 spores |
| *N. ironsii* (ERTm5) | 23,047 spores/μL | 27.4μL, ~ 6.32 x 10^5 spores | 63.8μL, ~ 14.70 x 10^5 spores |

Doses were calibrated based on the results of infectivity assays so that ERTm1 and ERTm5 inocula were comparably infective at a given dose.

**Table C: Number of hermaphrodite hosts per strain and treatment in the fitness assay.**

*Np* = *N. parisii*; *Ni* = *N. ironsii*. *low* = low dose; *high* = high dose.

| | | Treatment | | | | |
|---|---|---|---|---|---|---|
| **Host** | **Block** | *Control* | *Np low* | *Np high* | *Ni low* | *Ni high* |
| **CB4856** | 1 | 9 | 9 | 10 | 9 | 10 |
| **DL238** | 4 | 9 | 8 | 8 | 7 | 9 |
| **ECA1977** | 6 | 10 | 9 | 9 | 10 | 10 |
| **QX1791** | 5 | 10 | 10 | 10 | 9 | 10 |
| **ECA372** | 2 | 10 | 8 | 7 | 8 | 6 |
| **ECA705** | 4 | 10 | 10 | 10 | 10 | 10 |
| **ECA743** | 3 | 10 | 9 | 10 | 10 | 9 |
| **ECA812** | 3 | 9 | 10 | 9 | 10 | 10 |
| **QX1792** | 6 | 10 | 10 | 10 | 10 | 10 |
| **ECA2334** | 5 | 10 | 10 | 9 | 10 | 10 |
| **ECA347** | 1 | 9 | 10 | 10 | 10 | 10 |
| **ECA363** | 2 | 10 | 10 | 10 | 10 | 10 |
| **ECA724** | 2 | 9 | 10 | 9 | 10 | 9 |
| **ECA740** | 3 | 10 | 9 | 10 | 10 | 10 |
| **XZ1514** | 4 | 10 | 7 | 10 | 9 | 9 |
| **ECA1997** | 4 | 8 | 7 | 8 | 7 | 9 |
| **ECA730** | 2 | 10 | 10 | 10 | 10 | 9 |
| **ECA744** | 5 | 10 | 7 | 8 | 6 | 7 |
| **ECA746** | 6 | 10 | 10 | 9 | 10 | 9 |
| **N2*** | 1-6 | 26 | 30 | 30 | 29 | 30 |

*We included the strain N2 in each block to evaluate block effects. N2 was represented in full in block 1 (10 hosts per treatment), then with smaller sample sizes in the subsequent five blocks (4 hosts per treatment).

**Table D: Statistical analysis of total fecundity in control and exposed conditions across host strains**

A. Full model

| | |
|---|---|
| *Response* | Number of offspring |
| *Fixed effects* | Host strain*Condition |
| *Random effects* | Block and Plate ID |
| *Distribution* | Gaussian |

Condition = Control or Exposed, with Exposed encompassing all four exposed treatments.

B. Likelihood ratio test

| Model | Fixed effects | AIC | $\chi^2$ | df | *p* |
|---|---|---|---|---|---|
| 1 | Host strain*Condition | 10813 | | | |
| | | | 74.6 | 19 | <0.001 |
| 2 | Host strain + Condition | 10849 | | | |

C. Summary of best model

*Reference = CB4856, Control*

| Fixed Effect | Level | HostID | Coefficient ± SE | t value |
|---|---|---|---|---|
| Intercept | | | 141.9 ± 24.3 | 5.8 |
| Host strain | DL238 | 14 | 22.9 ± 34.2 | 0.7 |
| | ECA1977 | 16 | 17.9 ± 33.8 | 0.5 |
| | ECA1997 | 8 | -11.8 ± 23.7 | -0.5 |
| | ECA2334 | 13 | 2.5 ± 22.6 | 0.1 |
| | ECA347 | 15 | 21.3 ± 32.7 | 0.7 |
| | ECA363 | 7 | -7.4 ± 22.6 | -0.3 |
| | ECA372 | 10 | 15.7 ± 33.8 | 0.5 |
| | ECA705 | 5 | -18.7 ± 33.8 | -0.6 |
| | ECA724 | 6 | -8.1 ± 34.1 | -0.2 |
| | ECA730 | 4 | -15.1 ± 33.8 | -0.4 |
| | ECA740 | 3 | -48.5 ± 22.6 | -2.1 |
| | ECA743 | 12 | 16.2 ± 33.8 | 0.5 |
| | ECA744 | 19 | 61.1 ± 33.8 | 1.8 |
| | ECA746 | 17 | 19.6 ± 22.5 | 0.9 |
| | ECA812 | 2 | -61 ± 34.1 | -1.8 |
| | N2 | 20 | 84.8 ± 30.7 | 2.8 |
| | QX1791 | 18 | 29.0 ± 33.8 | 0.9 |
| | QX1792 | 11 | -3.9 ± 33.8 | -0.1 |
| | XZ1514 | 1 | -108.8 ± 33.8 | -3.2 |
| Condition | Exposed | | -91.1 ± 25.8 | -3.5 |
| Host strain:Condition | DL238:Exposed | 14 | 28.3 ± 36.6 | 0.8 |
| | ECA1977:Exposed | 16 | 11.7 ± 36.2 | 0.3 |
| | ECA1997:Exposed | 8 | 9.9 ± 24.1 | 0.4 |
| | ECA2334:Exposed | 13 | 6.7 ± 22.7 | 0.3 |
| | ECA347:Exposed | 15 | 0.7 ± 36.4 | 0.0 |
| | ECA363:Exposed | 7 | 27.0 ± 22.7 | 1.2 |
| | ECA372:Exposed | 10 | 62.2 ± 36.4 | 1.7 |
| | ECA705:Exposed | 5 | 106.4 ± 36.1 | 2.9 |
| | ECA724:Exposed | 6 | 60.5 ± 36.5 | 1.7 |
| | ECA730:Exposed | 4 | 28.0 ± 36.2 | 0.8 |
| | ECA740:Exposed | 3 | 56.4 ± 22.7 | 2.5 |
| | ECA743:Exposed | 12 | 37.9 ± 36.2 | 1.0 |
| | ECA744:Exposed | 19 | -20.0 ± 36.4 | -0.5 |
| | ECA746:Exposed | 17 | 17.0 ± 22.8 | 0.7 |
| | ECA812:Exposed | 2 | 68.1 ± 36.5 | 1.9 |
| | N2:Exposed | 20 | -14.3 ± 33.9 | -0.4 |
| | QX1791:Exposed | 18 | -12.4 ± 36.2 | -0.3 |
| | QX1792:Exposed | 11 | 62.3 ± 36.1 | 1.7 |

| | | | |
|---|---|---|---|
| XZ1514:Exposed | 1 | 72.4 ± 36.2 | 2.0 |

**Table E: Statistical analysis of total fecundity in control and exposed conditions across host group**

A. Full model

| | |
|---|---|
| *Response* | Number of offspring |
| *Fixed effects* | Host group * Condition |
| *Random effects* | Host strain and Plate ID |
| *Distribution* | Gaussian |

Condition = Control or Exposed, with Exposed encompassing all four exposed treatments.
Block is not included as a random effect, because N2 is excluded from these analyses. N2 is not part of a Hawaiian group.

B. Likelihood ratio test

| Model | Fixed effects | AIC | $\chi^2$ | df | *p* |
|---|---|---|---|---|---|
| 1 | Host group*Condition | 9242 | | | |
| | | | 41.6 | 3 | <0.001 |
| 2 | Host group + Condition | 9277 | | | |

C. Summary of best model

*Reference = Divergent, Control*

| Fixed Effect | Level | Coefficient ± SE | t value |
|---|---|---|---|
| Intercept | | 109.4 ± 15.5 | 7.1 |
| Host group | Invaded | 16.9 ± 20.6 | 0.8 |
| | Low | 54.7 ± 20.9 | 2.6 |
| | Volcano | 44.4 ± 20.9 | 2.1 |
| Condition | Exposed | -48.8 ± 10.9 | -4.5 |
| Host group:Condition | Invaded:Exposed | 27.2 ± 11.2 | 2.4 |
| | Low:Exposed | -35.7 ± 10.4 | -3.4 |
| | Volcano:Exposed | -28.3 ± 10.4 | -2.7 |

### Table F: Statistical analysis of total fecundity across treatments

#### A. Full model

| | |
|---|---|
| *Response* | Number of offspring |
| *Fixed effects* | Host strain * Treatment |
| *Random effects* | Block |
| *Distribution* | Gaussian |

Treatment = Control, *N. parisii* at low dose, *N. parisii* at high dose, *N. ironsii* at low dose, *N. ironsii* at high dose

#### B. Likelihood ratio test

| Model | Fixed effects | AIC | $\chi^2$ | df | *p* |
|---|---|---|---|---|---|
| 1 | Host strain * Treatment | 10820 | | | |
| | | | 161.9 | 76 | <0.001 |
| 2 | Host strain + Treatment | 10830 | | | |

#### C. Summary of model 2 to highlight treatment main effects

*Reference = Control treatment, CB4856*

| Fixed Effect | Level | HostID | Coefficient ± SE | t value |
|---|---|---|---|---|
| Intercept | | | 119.2 ± 10.7 | 11.2 |
| Treatment | *N. parisii*, low dose | | -42.5 ± 4.6 | -9.3 |
| | *N. parisii*, high dose | | -64.9 ± 4.5 | -14.3 |
| | *N. ironii*, low dose | | -63.6 ± 4.5 | -14.0 |
| | *N. ironsii*, high dose | | -84.5 ± 4.5 | -18.6 |
| Host strain | DL238 | 14 | 46.9 ± 14.1 | 3.3 |
| | ECA1977 | 16 | 28.0 ± 13.7 | 2.0 |
| | ECA1997 | 8 | -2.6 ± 14.2 | -0.2 |
| | ECA2334 | 13 | 9.9 ± 13.8 | 0.7 |
| | ECA347 | 15 | 21.6 ± 9.4 | 2.3 |
| | ECA363 | 7 | 15.1 ± 13.7 | 1.1 |
| | ECA372 | 10 | 64.3 ± 14.2 | 4.5 |
| | ECA705 | 5 | 67.3 ± 13.8 | 4.9 |
| | ECA724 | 6 | 41.3 ± 13.8 | 3.0 |
| | ECA730 | 4 | 8.1 ± 13.8 | 0.6 |
| | ECA740 | 3 | -2.2 ± 13.8 | -0.2 |
| | ECA743 | 12 | 47.7 ± 13.8 | 3.5 |
| | ECA744 | 19 | 50.2 ± 14.2 | 3.5 |
| | ECA746 | 17 | 33.8 ± 13.7 | 2.5 |
| | ECA812 | 2 | -4.7 ± 13.8 | -0.3 |
| | N2 | 20 | 73.3 ± 9.0 | 8.2 |
| | QX1791 | 18 | 21.2 ± 13.8 | 1.5 |
| | QX1792 | 11 | 46.6 ± 13.6 | 3.4 |
| | XZ1514 | 1 | -50.7 ± 14.0 | -3.6 |

**Table G: Statistical analysis of change in total fecundity from control for each exposure treatment**

A. Representative full model

| | |
|---|---|
| *Response* | Number of offspring |
| *Fixed effects* | Host strain * Treatment |
| *Random effects* | Block |
| *Distribution* | Gaussian |

Treatment = Control and EITHER *N. parisii* at low dose, *N. parisii* at high dose, *N. ironsii* at low dose, OR *N. ironsii* at high dose

B. Low dose of *N. parisii* vs. control, likelihood ratio test

| **Model** | **Fixed effects** | **AIC** | $\chi^2$ | **df** | ***p*** |
|---|---|---|---|---|---|
| 1 | Host strain*Treatment | 4472 | | | |
| | | | 49.6 | 19 | <0.001 |
| 2 | Host strain + Treatment | 4461 | | | |

C. High dose of *N. parisii* vs. control, likelihood ratio test

| **Model** | **Fixed effects** | **AIC** | $\chi^2$ | **df** | ***p*** |
|---|---|---|---|---|---|
| 1 | Host strain*Treatment | 4392 | | | |
| | | | 78.1 | 19 | <0.001 |
| 2 | Host strain + Treatment | 4432 | | | |

D. Low dose of *N. ironsii* vs. control, likelihood ratio test

| **Model** | **Fixed effects** | **AIC** | $\chi^2$ | **df** | ***p*** |
|---|---|---|---|---|---|
| 1 | Host strain*Treatment | 4404 | | | |
| | | | 52.2 | 19 | <0.001 |
| 2 | Host strain + Treatment | 4418 | | | |

E. High dose of *N. ironsii* vs. control, likelihood ratio test

| **Model** | **Fixed effects** | **AIC** | $\chi^2$ | **df** | ***p*** |
|---|---|---|---|---|---|
| 1 | Host strain*Treatment | 4405 | | | |
| | | | 68.9 | 19 | <0.001 |
| 2 | Host strain + Treatment | 4436 | | | |

### Table H: Statistical analysis of total fecundity across exposed treatments

#### A. Full model

| | |
|---|---|
| *Response* | Number of offspring |
| *Fixed effects* | Host strain * Exposure Treatment |
| *Random effects* | Block |
| *Distribution* | Gaussian |

Exposure treatment = *N. parisii* at low dose, *N. parisii* at high dose, *N. ironsii* at low dose, *N. ironsii* at high dose (**Control excluded**)

#### B. Likelihood ratio test

| Model | Fixed effects | AIC | $\chi^2$ | df | *p* |
|---|---|---|---|---|---|
| 1 | Host strain * Treatment | 8530 | | | |
| | | | 69.6 | 57 | 0.123 |
| 2 | Host strain + Treatment | 8486 | | | |

#### C. Summary of best model

*Reference* = N. parisii, *low dose treatment, CB4856*

| Fixed Effect | Level | HostID | Coefficient ± SE | t value |
|---|---|---|---|---|
| Intercept | | | 69.8 ± 11.4 | 6.1 |
| Treatment | *N. parisii*, high dose | | -22.4 ± 4.2 | -5.4 |
| | *N. ironii*, low dose | | -21.4 ± 4.2 | -5.1 |
| | *N. ironsii*, high dose | | -42.0 ± 4.2 | -10.1 |
| Host strain | DL238 | 14 | 55.1 ± 14.8 | 3.7 |
| | ECA1977 | 16 | 24.5 ± 14.4 | 1.7 |
| | ECA1997 | 8 | 1.8 ± 14.8 | 0.1 |
| | ECA2334 | 13 | 13.8 ± 14.3 | 1.0 |
| | ECA347 | 15 | 21.9 ± 9.5 | 2.3 |
| | ECA363 | 7 | 27.2 ± 14.3 | 1.9 |
| | ECA372 | 10 | 85.4 ± 14.9 | 5.7 |
| | ECA705 | 5 | 91 ± 14.4 | 6.3 |
| | ECA724 | 6 | 59.8 ± 14.4 | 4.2 |
| | ECA730 | 4 | 20.5 ± 14.3 | 1.4 |
| | ECA740 | 3 | 12.5 ± 14.3 | 0.9 |
| | ECA743 | 12 | 58.5 ± 14.4 | 4.1 |
| | ECA744 | 19 | 45.9 ± 14.9 | 3.1 |
| | ECA746 | 17 | 31.4 ± 14.4 | 2.2 |
| | ECA812 | 2 | 11.6 ± 14.3 | 0.8 |
| | N2 | 20 | 72.4 ± 9.2 | 7.9 |
| | QX1791 | 18 | 21.2 ± 14.3 | 1.5 |
| | QX1792 | 11 | 53.3 ± 14.3 | 3.7 |
| | XZ1514 | 1 | -32.8 ± 14.6 | -2.2 |

### Table I: Correlation[1] of defense[2] across exposure treatments

| $\rho$ | *N. parisii,* low dose | *N. parisii,* high dose | *N. ironsii,* low dose | *N. ironsii,* high dose |
|---|---|---|---|---|
| *N. parisii,* low dose | 1.00 | | | |
| *N. parisii,* high dose | 0.65* | 1.00 | | |
| *N. ironsii,* low dose | 0.60* | 0.51 | 1.00 | |
| *N. ironsii,* high dose | 0.80* | 0.72* | 0.63* | 1.00 |

[1] Spearman's rank correlation coefficient $\rho$ was calculated because defense estimates were not consistently normally distributed.

[2] Defense calculated for each host strain as the mean number of offspring per host in an exposed treatment, divided by the mean number per host in the control treatment.

[*] denotes p-value is less than alpha value of 0.008, after Bonferroni correction for six tests.

**Table J: Statistical analysis of daily fecundity by host strain in the control treatment**

A. Full model

| | |
|---|---|
| *Response* | Number of offspring per day |
| *Fixed effects* | Host strain*Day |
| *Random effects* | Individual ID, Block |
| *Distribution* | Gaussian |

B. Likelihood ratio test

| Model | Fixed effects | AIC | $\chi^2$ | df | *p* |
|---|---|---|---|---|---|
| 1 | Host strain * Day | 9237 | | | |
| | | | 425.1 | 76 | <0.001 |
| 2 | Host strain + Day | 9510 | | | |

Part C excluded due to its length

**Table K: Statistical analysis of daily fecundity by host group in the control treatment**

A. Full model

| | |
|---|---|
| *Response* | Number of offspring per day |
| *Fixed effects* | Host group*Day |
| *Random effects* | Host strain, Individual ID |
| *Distribution* | Gaussian |

Block is not included as a random effect, because N2 is excluded from these analyses. N2 is not part of a Hawaiian group

B. Likelihood ratio test

| Model | Fixed effects | AIC | $\chi^2$ | df | *p* |
|---|---|---|---|---|---|
| 1 | Host group * Day | 8069 | | | |
| | | | 103.1 | 12 | <0.001 |
| 2 | Host group + Day | 8184 | | | |

C. Summary of best model

*Reference = Day 1, Divergent group*

| Fixed Effect | Level | Coefficient ± SE | t value |
|---|---|---|---|
| Intercept | | 7.2 ± 4.0 | 1.8 |
| Day | 2 | 54.8 ± 3.8 | 14.4 |
| | 3 | 28.5 ± 3.8 | 7.5 |
| | 4 | 0.50 ± 3.8 | 0.1 |
| | 5 | -5.6 ± 3.8 | -1.5 |
| Host group | Invaded | 5.3 ± 5.9 | 0.9 |
| | Low | 6.2 ± 6.3 | 1.0 |
| | Volcano | 5.9 ± 6.3 | 0.9 |
| Host group:Day | Invaded:2 | 13.7 ± 5.7 | 2.4 |
| | Invaded:3 | -8.8 ± 5.7 | -1.6 |
| | Invaded:4 | -7.8 ± 5.7 | -1.4 |
| | Invaded:5 | -5.3 ± 5.7 | -0.9 |
| | Low:2 | -2.2 ± 6.0 | -0.4 |
| | Low:3 | 27.1 ± 6.0 | 4.5 |
| | Low:4 | -0.3 ± 6.0 | 0.0 |
| | Low:5 | -6.0 ± 6.0 | -1.0 |
| | Volcano:2 | 2.0 ± 6.0 | 0.3 |
| | Volcano:3 | 22.1 ± 6.0 | 3.7 |
| | Volcano:4 | -4.4 ± 6.0 | -0.7 |
| | Volcano:5 | -6.6 ± 6.0 | -1.1 |

### Table L: Statistical analysis of daily fecundity in control and exposed conditions

A. Full model

| | |
|---|---|
| *Response* | Number of offspring per day |
| *Fixed effects* | Condition*Day |
| *Random effects* | Individual ID, Host strain, Block, Plate ID |
| *Distribution* | Gaussian |

Condition = Control or Exposed, with Exposed encompassing all four exposure treatments.

B. Likelihood ratio test

| Model | Fixed effects | AIC | $\chi^2$ | df | *p* |
|---|---|---|---|---|---|
| 1 | Condition * Day | 43914 | | | |
| | | | 444.0 | 4 | <0.001 |
| 2 | Condition + Day | 44350 | | | |

C. Summary of best model

*Reference = Control, Day 2*

| Fixed Effect | Level | Coefficient ± SE | t value |
|---|---|---|---|
| Intercept | | 75.8 ± 2.8 | 26.9 |
| Day | 1 | -64.7 ± 1.8 | -35.5 |
| | 3 | -24.8 ± 1.8 | -13.6 |
| | 4 | -67.1 ± 1.8 | -36.8 |
| | 5 | -74.1 ± 1.8 | -40.7 |
| Condition | Exposed | -27.5 ± 2.3 | -11.9 |
| Condition:Day | Exposed:1 | 26.7 ± 2.0 | 13.1 |
| | Exposed:3 | -4.9 ± 2.0 | -2.4 |
| | Exposed:4 | 23.8 ± 2.0 | 11.6 |
| | Exposed:5 | 26.9 ± 2.0 | 13.2 |

### Table M: Linear regression of reproductive timing and defense against parasites

| Fixed Effect | Coefficient ± SE | t value | *p* |
|---|---|---|---|
| Intercept | 0.12 ± 0.16 | 0.73 | 0.477 |
| Fraction early | 0.78 ± 0.27 | 2.92 | 0.009 |

$R^2$ = 0.321
Adj. $R^2$ = 0.284

Defense = mean total offspring per host when exposed/mean total offspring per host in control, for each host strain
Reproductive timing = mean fraction of total offspring per host made on days 1 and 2 of reproduction, for each host strain, in the control treatment

### Table N: Linear regression of fecundity and defense against parasites

| Fixed Effect | Coefficient ± SE | t value | *p* |
|---|---|---|---|
| Intercept | 0.60 ± 0.15 | 4.11 | <0.001 |
| Fecundity | -0.00 ± 0.00 | -0.21 | 0.839 |

$R^2$ = 0.002
Adj. $R^2$ = -0.053

Defense = mean total offspring per host when exposed/mean total offspring per host in control, for each host strain
Fecundity = mean total offspring per host in control treatment, for each host strain

**Table O: Replication structure for resistance assays.** Columns are the replicate populations assayed for resistance, and numbers indicate the number of hermaphrodite hosts measured per replicate.

| | 48 hours | | | | 72 hours | | | |
|---|---|---|---|---|---|---|---|---|
| **Strain/isotype** | **1** | **2** | **3** | **4** | **1** | **2** | **3** | **4** |
| **CB4856** | 132 | 50 | 50 | | 35 | 35 | 35 | 35 |
| **DL238** | 120 | 51 | 35 | | | | | |
| **ECA1977** | 51 | 50 | 34 | | 35 | 35 | 35 | |
| **QX1791** | 49 | 49 | 35 | | | | | |
| **ECA372** | 50 | 49 | 35 | | | | | |
| **ECA705** | 51 | 49 | 51 | | 35 | 35 | 35 | 35 |
| **ECA743** | 35 | 49 | 50 | | | | | |
| **ECA812** | 49 | 49 | 31 | | 32 | 34 | 34 | 34 |
| **QX1792** | 49 | 49 | 35 | | | | | |
| **ECA2334** | 50 | 50 | 50 | | 35 | 36 | 35 | 37 |
| **ECA347** | 47 | 41 | 35 | | | | | |
| **ECA363** | 49 | 50 | 34 | | 35 | 34 | 35 | |
| **ECA724** | 45 | 35 | 50 | | 35 | 35 | 35 | 36 |
| **ECA740** | 49 | 50 | 50 | | | | | |
| **XZ1514** | 49 | 55 | 50 | | | | | |
| **ECA1997** | 49 | 49 | 36 | | 36 | 35 | | |
| **ECA730** | 49 | 50 | 35 | | | | | |
| **ECA744** | 31 | 55 | 50 | 33 | 35 | 35 | 36 | |
| **ECA746** | 53 | 51 | 50 | | | | | |
| **N2** | 85 | 49 | 35 | | 33 | 35 | 35 | |

**Table P: Statistical analysis of infection prevalence and load at 48 hours.**

A. Full model

| | |
|---|---|
| *Response* | Fraction of body area fluorescent |
| *Fixed effects* | Host strain |
| *Random effects* | Replicate, Block |
| *Zero-inflation* | Host strain |
| *Distribution* | Beta |

Fluorescence indicates presence of *N. parisii*

B. Likelihood ratio test

*Zero-inflation model: do host strains vary in infection prevalence?*

| Model | Zero-inflation term | AIC | $\chi^2$ | df | *p* |
|---|---|---|---|---|---|
| 1 | Host strain | -12017 | | | |
| | | | 84.5 | 19 | <0.001 |
| 2 | Intercept-only | -11970 | | | |

*Conditional model: do host strains vary in infection load?*

| Model | Fixed effect | AIC | $\chi^2$ | df | *p* |
|---|---|---|---|---|---|
| 1 | Host strain | -12017 | | | |
| | | | 51.2 | 19 | <0.001 |
| 2 | Intercept-only | -12003 | | | |

C. Summary of best model

*Reference = CB4856*

*Zero-inflation model: do host strains vary in infection prevalence?*

| Term | Level | HostID | Coefficient ± SE | Odds ratio [95%CI] | z value |
|---|---|---|---|---|---|
| Intercept | | | -1.37 ± 0.16 | 0.25 [0.18,0.35] | -8.4 |
| Host strain | DL238 | 14 | 0.21 ± 0.23 | 1.23 [0.78,1.93] | 0.9 |
| | ECA1977 | 16 | 0.58 ± 0.25 | 1.78 [1.09,2.89] | 2.3 |
| | ECA1997 | 8 | -0.20 ± 0.28 | 0.82 [0.47,1.42] | -0.7 |
| | ECA2334 | 13 | 0.59 ± 0.24 | 1.80 [1.12,2.88] | 2.4 |
| | ECA347 | 15 | -0.68 ± 0.33 | 0.51 [0.27,0.96] | -2.1 |
| | ECA363 | 7 | -0.14 ± 0.28 | 0.87 [0.50,1.50] | -0.5 |
| | ECA372 | 10 | -0.31 ± 0.29 | 0.73 [0.42,1.29] | -1.1 |
| | ECA705 | 5 | 0.28 ± 0.25 | 1.32 [0.81,2.16] | 1.1 |
| | ECA724 | 6 | 0.17 ± 0.26 | 1.18 [0.70,1.98] | 0.6 |
| | ECA730 | 4 | -0.15 ± 0.28 | 0.86 [0.50,1.48] | -0.5 |
| | ECA740 | 3 | -0.05 ± 0.26 | 0.95 [0.57,1.59] | -0.2 |
| | ECA743 | 12 | 0.08 ± 0.27 | 1.09 [0.65,1.83] | 0.3 |
| | ECA744 | 19 | -1.93 ± 0.45 | 0.14 [0.06,0.35] | -4.3 |
| | ECA746 | 17 | -0.13 ± 0.27 | 0.87 [0.52,1.47] | -0.5 |
| | ECA812 | 2 | -0.16 ± 0.28 | 0.85 [0.49,1.48] | -0.6 |
| | N2 | 20 | -0.43 ± 0.27 | 0.65 [0.38,1.12] | -1.6 |
| | QX1791 | 18 | 0.22 ± 0.26 | 1.25 [0.75,2.08] | 0.8 |
| | QX1792 | 11 | -0.42 ± 0.30 | 0.66 [0.37,1.17] | -1.4 |
| | XZ1514 | 1 | 0.18 ± 0.25 | 1.20 [0.73,1.96] | 0.7 |

*Conditional model: do host strains vary in infection load?*

| Fixed Effect | Level | HostID | Coefficient ± SE | Odds ratio [95%CI] | z value |
|---|---|---|---|---|---|
| Intercept | | | -3.80 ± 0.11 | 0.02 [0.02,0.03] | -34.1 |
| Host strain | DL238 | 14 | -0.04 ± 0.11 | 0.96 [0.77,1.19] | -0.4 |
| | ECA1977 | 16 | 0.28 ± 0.13 | 1.32 [1.03,1.69] | 2.2 |
| | ECA1997 | 8 | -0.31 ± 0.14 | 0.73 [0.56,0.96] | -2.3 |
| | ECA2334 | 13 | 0.01 ± 0.15 | 1.01 [0.75,1.35] | 0.1 |
| | ECA347 | 15 | -0.23 ± 0.14 | 0.80 [0.60,1.05] | -1.6 |
| | ECA363 | 7 | 0.22 ± 0.14 | 1.25 [0.95,1.64] | 1.6 |
| | ECA372 | 10 | 0.11 ± 0.14 | 1.12 [0.84,1.48] | 0.8 |
| | ECA705 | 5 | -0.13 ± 0.14 | 0.88 [0.68,1.15] | -0.9 |
| | ECA724 | 6 | 0.07 ± 0.14 | 1.07 [0.81,1.42] | 0.5 |
| | ECA730 | 4 | 0.00 ± 0.14 | 1 .00[0.75,1.32] | 0.0 |
| | ECA740 | 3 | 0.10 ± 0.14 | 1.11 [0.85,1.45] | 0.8 |
| | ECA743 | 12 | -0.30 ± 0.15 | 0.74 [0.56,0.98] | -2.1 |
| | ECA744 | 19 | 0.33 ± 0.13 | 1.40 [1.08,1.80] | 2.6 |
| | ECA746 | 17 | 0.22 ± 0.14 | 1.25 [0.95,1.64] | 1.6 |
| | ECA812 | 2 | 0.03 ± 0.14 | 1.03 [0.78,1.36] | 0.2 |
| | N2 | 20 | 0.39 ± 0.14 | 1.48 [1.12,1.94] | 2.8 |
| | QX1791 | 18 | 0.25 ± 0.13 | 1.29 [1.00,1.66] | 2.0 |
| | QX1792 | 11 | 0.14 ± 0.13 | 1.15 [0.88,1.49] | 1.0 |
| | XZ1514 | 1 | 0.00 ± 0.14 | 1 .00[0.76,1.30] | 0.0 |

**Table Q: Statistical analysis of infection prevalence and load by group at 48 hours**.

A. Full model

| | |
|---|---|
| *Response* | Fraction of body area fluorescent |
| *Fixed effects* | Host group |
| *Random effects* | Host strain, Replicate, Block |
| *Zero-inflation* | Host group |
| *Distribution* | Beta |

Fluorescence indicates presence of *N. parisii.* N2 is excluded from these analyses because it is not part of a Hawaiian group

B. Likelihood ratio test

*Zero-inflation model: do host groups vary in infection prevalence?*

| Model | Zero-inflation term | AIC | $\chi^2$ | df | *p* |
|---|---|---|---|---|---|
| 1 | Host group | -11370 | | | |
| | | | 24.5 | 3 | <0.001 |
| 2 | Intercept-only | -11351 | | | |

*Conditional model: do host groups vary in infection load?*

| Model | Fixed effect | AIC | $\chi^2$ | df | *p* |
|---|---|---|---|---|---|
| 1 | Host group | -11370 | | | |
| | | | 1.8 | 3 | 0.614 |
| 2 | Intercept-only | -11374 | | | |

C. Summary of best model

*Reference = Divergent group*

*Zero-inflation model: do host groups vary in infection prevalence?*

| Term | Level | Coefficient ± SE | Odds ratio [95%CI] | z value |
|---|---|---|---|---|
| Intercept | | -1.30 ± 0.08 | 0.27 [0.23,0.32] | -15.4 |
| Host group | Invaded | -0.15 ± 0.13 | 0.86 [0.67,1.11] | -1.1 |
| | Low | 0.15 ± 0.12 | 1.16 [0.91,1.47] | 1.2 |
| | Volcano | -0.54 ± 0.15 | 0.58 [0.44,0.77] | -3.7 |

*Conditional model: do host groups vary in infection load?*

| Fixed Effect | Level | Coefficient ± SE | Odds ratio [95%CI] | z value |
|---|---|---|---|---|
| Intercept | | -3.76 ± 0.09 | 0.02 [0.02,0.03] | -41.0 |
| Host group | Invaded | -0.09 ± 0.1 | 0.92 [0.75,1.12] | -0.8 |
| | Low | 0.06 ± 0.12 | 1.06 [0.84,1.34] | 0.5 |
| | Volcano | 0.03 ± 0.11 | 1.03 [0.84,1.27] | 0.3 |

**Table R: Statistical analysis of infection load at 72 hours**.

A. Full model

| | |
|---|---|
| *Response* | Fraction of body area fluorescent |
| *Fixed effects* | Host strain |
| *Random effects* | Replicate, Block |
| *Distribution* | Beta |

Fluorescence indicates presence of *N. parisii.* Only 5 out of 1,187 had no sign of infection (i.e., no fluorescence), so these hosts are excluded to specifically evaluate variation in load of infected hosts.

B. Likelihood ratio test

| **Model** | **Fixed effect** | **AIC** | $\chi^2$ | **df** | ***p*** |
|---|---|---|---|---|---|
| 1 | Host strain | -2983 | | | |
| | | | 30.3 | 9 | <0.001 |
| 2 | Intercept-only | -2971 | | | |

C. Summary of best model

*Reference = CB4856*

| **Fixed Effect** | **Level** | **HostID** | **Coefficient ± SE** | **Odds ratio [95%CI]** | **z value** |
|---|---|---|---|---|---|
| Intercept | | | -1.72 ± 0.19 | 0.18 [0.12,0.26] | -9.1 |
| Host strain | ECA1977 | 16 | -0.35 ± 0.12 | 0.70 [0.56,0.88] | -3.0 |
| | ECA1997 | 8 | -0.24 ± 0.15 | 0.78 [0.59,1.04] | -1.7 |
| | ECA2334 | 13 | -0.72 ± 0.12 | 0.48 [0.38,0.62] | -5.8 |
| | ECA363 | 7 | -0.18 ± 0.13 | 0.84 [0.65,1.08] | -1.4 |
| | ECA705 | 5 | -0.55 ± 0.12 | 0.58 [0.46,0.73] | -4.7 |
| | ECA724 | 6 | -0.45 ± 0.11 | 0.64 [0.51,0.80] | -3.9 |
| | ECA744 | 19 | -0.48 ± 0.13 | 0.62 [0.48,0.79] | -3.8 |
| | ECA812 | 2 | -0.58 ± 0.11 | 0.56 [0.45,0.70] | -5.2 |
| | N2 | 20 | -0.41 ± 0.13 | 0.66 [0.51,0.86] | -3.1 |

**Table T: Statistical analysis of infection load by group at 72 hours**.

A. Full model

| | |
|---|---|
| *Response* | Fraction of body area fluorescent |
| *Fixed effects* | Host group |
| *Random effects* | Host strain, Replicate, Block |
| *Distribution* | Beta |

Fluorescence indicates presence of *N. parisii.* Only 5 out of 1,187 had no sign of infection (i.e., no fluorescence), so these hosts are excluded to specifically evaluate variation in load of infected hosts. N2 is excluded from these analyses because it is not part of a Hawaiian group

B. Likelihood ratio test

| Model | Fixed effect | AIC | $\chi^2$ | df | *p* |
|---|---|---|---|---|---|
| 1 | Host group | -2696 | | | |
| | | | 4.5 | 3 | 0.216 |
| 2 | Intercept-only | -2697 | | | |

**Table U: Linear regression of resistance and defense against parasites**

A. 48 hours – 20 strains

| Fixed Effect | Coefficient ± SE | t value | *p* |
|---|---|---|---|
| Intercept | 0.76 ± 0.32 | 2.40 | 0.028 |
| Load | -0.02 ± 0.13 | -0.16 | 0.878 |

$R^2$ = 0.001
Adj. $R^2$ = -0.054

B. 72 hours – 10 strains

| Fixed Effect | Coefficient ± SE | t value | *p* |
|---|---|---|---|
| Intercept | 1.18 ± 0.43 | 2.74 | 0.026 |
| Load | -0.04 ± 0.03 | -1.10 | 0.302 |

$R^2$ = 0.132
Adj. $R^2$ = 0.024

Defense = mean total offspring per host when exposed to a low dose of ERTm1/mean total offspring per host in control, for each host strain
Resistance = mean predicted infection load, or percent body area infected, at 48 hours for each host strain. Predictions derived from the statistical model in Table S16, conditional component, to account for replicate and block effects.

### S2 TEXT – SUPPLEMENTAL SURVIVAL ANALYSIS

The fitness assay provided data on both survival and fecundity of hosts. Prior studies have demonstrated that the primary fitness effect of *Nematocida* infection is on host reproduction, not survival [1,2], and this was borne out in our dataset. We therefore focused on fecundity effects in the main text. Fecundity analyses included data from hosts that died over the course of the assay, unless it was due to experimental error, so fecundity data accounted for the negative effect of mortality on lifetime fecundity.

To specifically evaluate the effect of parasite exposure on survival, we fit a Cox proportional hazards model using the package *survival* [3]. We first included parasite exposure (yes or no) as a predictor of survival, comparing control hosts to hosts in all exposed treatments. In a second model, we included treatment as a predictor, comparing hosts across all five treatments. We excluded males from these analyses.

Parasite exposure substantially reduced survival during the fitness assay (hazard ratio = 5.28, $z = 8.85$, $p < 0.001$). In the absence of parasites, 14.3% of individuals died over the course of the assay (n = 30/209), while in the presence of parasites, 62.9% of individuals died (n = 515/819). Mortality was elevated in all exposure treatments, but it was relatively low under a low dose of *N. parisii* (43.4% died, hazard ratio = 3.73, $z = 5.75$, $p < 0.001$) and relatively high with a high dose of *N. ironsii* (80.1% died, hazard ratio = 6.94, $z = 9.75$, $p < 0.001$). Most exposed individuals died in the final day of reproduction: 21% of deaths occurred on the fourth day (n = 109/515), and 71% occurred on the fifth and final day (n = 367/515). Deaths thus primarily occurred after the bulk of reproduction: 91.8% of offspring had been produced before the fourth day of reproduction, and 98.2% had been produced before the final day (S7 Fig.). We accordingly view parasite-mediated reductions in survival as having relatively minor consequences for host fitness.

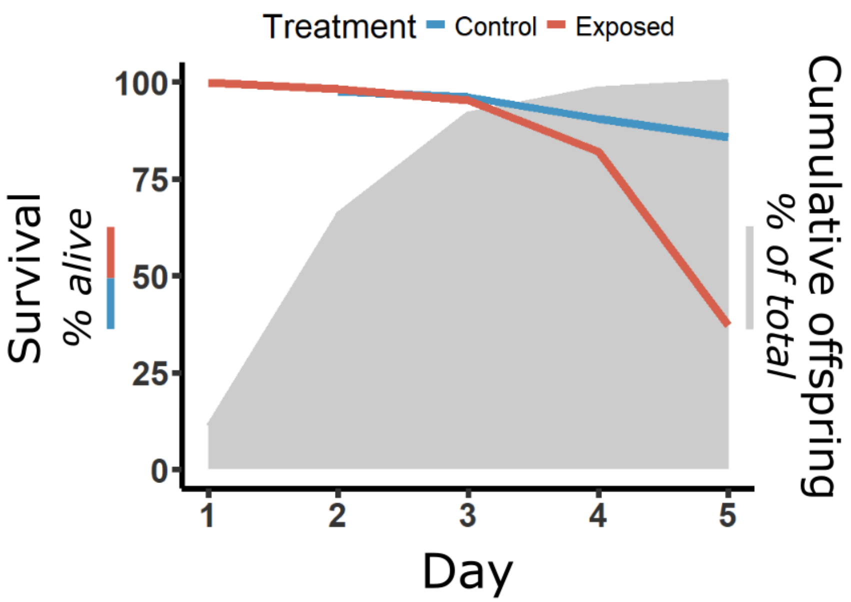


**S7 Fig: Parasite exposure reduces survival, but mortality occurs after reproduction.** Lines show the percentage of hosts alive during the fitness assay, in control (blue) and exposed (red) conditions. The Control group represents data from 209 hosts across 20 strains, while the Exposed group represents data from 819 hosts from 20 strains and four exposure treatments. Gray shaded area shows the cumulative offspring production through time across all hosts; this is represented as the cumulative percent of total offspring produced after a given day of reproduction.

### SUPPLEMENTAL FIGURES

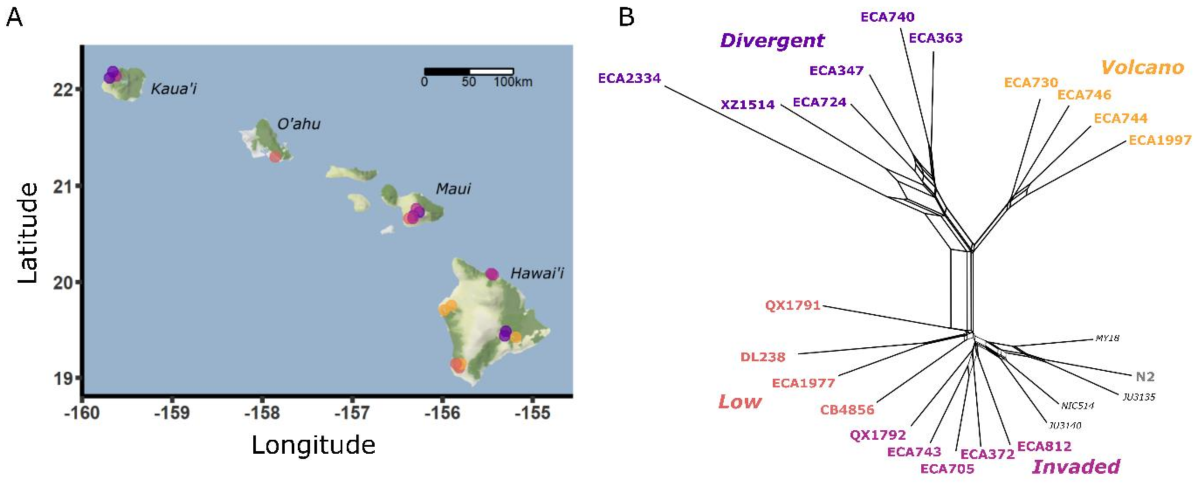


**S1 Fig: Host strains included in this study. A** shows the sampling locations of the 19 Hawaiian strains. Points are jittered to reduce overlap. The 20th strain, N2, is not from Hawaii and is not shown. The map was generated using the package *ggmap* [108]. The map tiles are by © Stamen Design, © Stadia Maps, and © OpenMapTiles under CC BY 4.0. Data by OpenStreetMap, under ODbL. **B** shows a neighbor-joining network of the 20 strains included in this study, plus four non-Hawaiian strains (in italics) that were not included in this study but are shown for added phylogenetic context. Colors of points in A and taxa in B correspond to the four relatedness clusters (i.e., group: Divergent, Volcano, Invaded, and Low). The neighbor-joining network was generated using [47]'s Figure 5 VCF dataset as the base variant call set. Three strains not included in that dataset (ECA1997, ECA1977, ECA2334) were downloaded from CaeNDR as VCF variant data and filtered to retain comparable sites. The additional strains were harmonized with [47]'s call set to generate a combined SNP matrix for the selected taxa. The merged VCF was converted to a NEXUS-formatted SNP alignment using vcf2phylip.py and then used to generate the neighbor-joining network in SplitsTree4 [109]. Table A in S1 Text provides further details on the host strains.

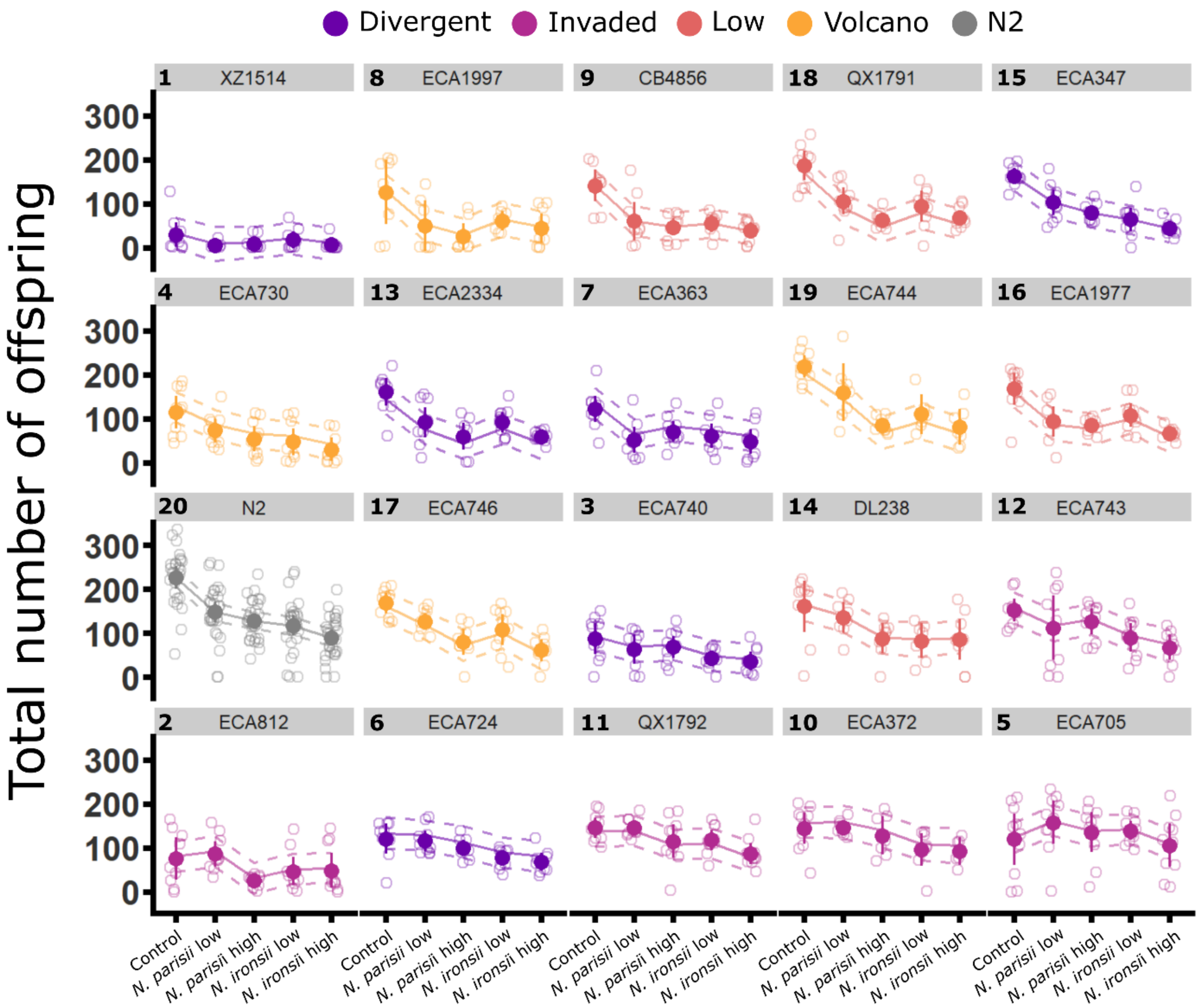


**S2 Fig: Fecundity of host strains across treatments.** The total number of offspring per host is shown for each of the five treatments, faceted by host strain. Points show raw data for total fecundity, while the lines show estimated marginal means from the full linear mixed effects model in Table F in S1 Text. For the raw data, unfilled points show the total number of offspring for individual hermaphrodites, and filled points show the mean. Error bars show 95% confidence intervals. For model estimates, solid lines indicate the mean and dashed lines the 95% confidence interval. Host strains are arrayed from top left to bottom right in order of increasing overall defense against *Nematocida*, as shown in Fig. 2a, and they are colored according to host group, with the host strain N2 in gray. Host strains are indicated by both their strain name and numeric ID.

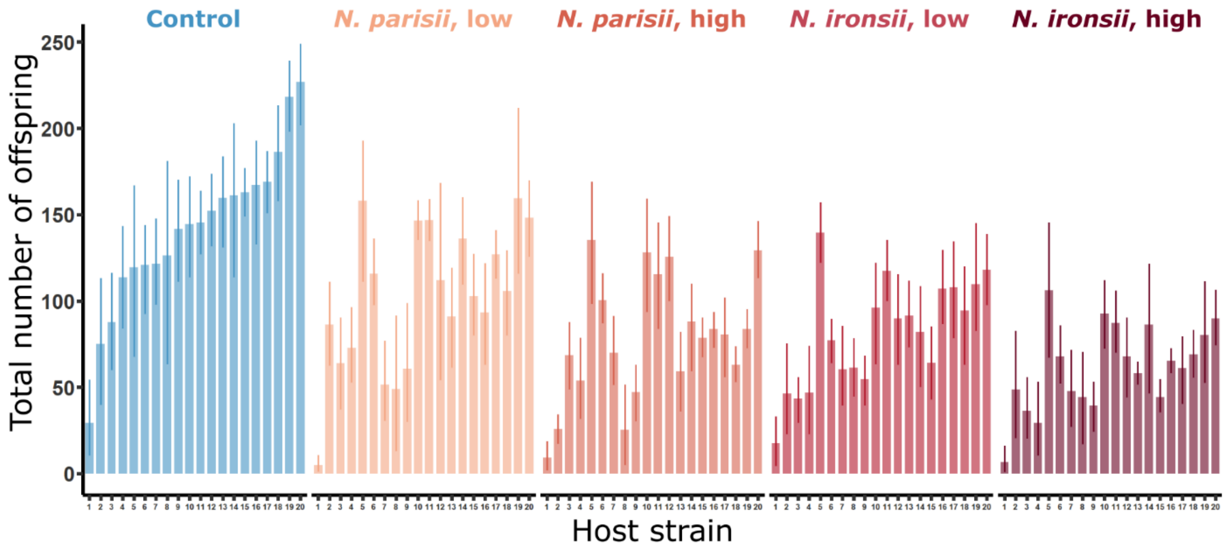


**S3 Fig: Variation in fecundity within treatments.** Bars show the mean number of offspring per host, with 95% confidence intervals, for each host strain. Within a treatment, host strains are ordered from left to right by increasing fecundity in baseline control conditions. Host strains are indicated by their numeric ID (Table A in S1 Text, Fig. 1).

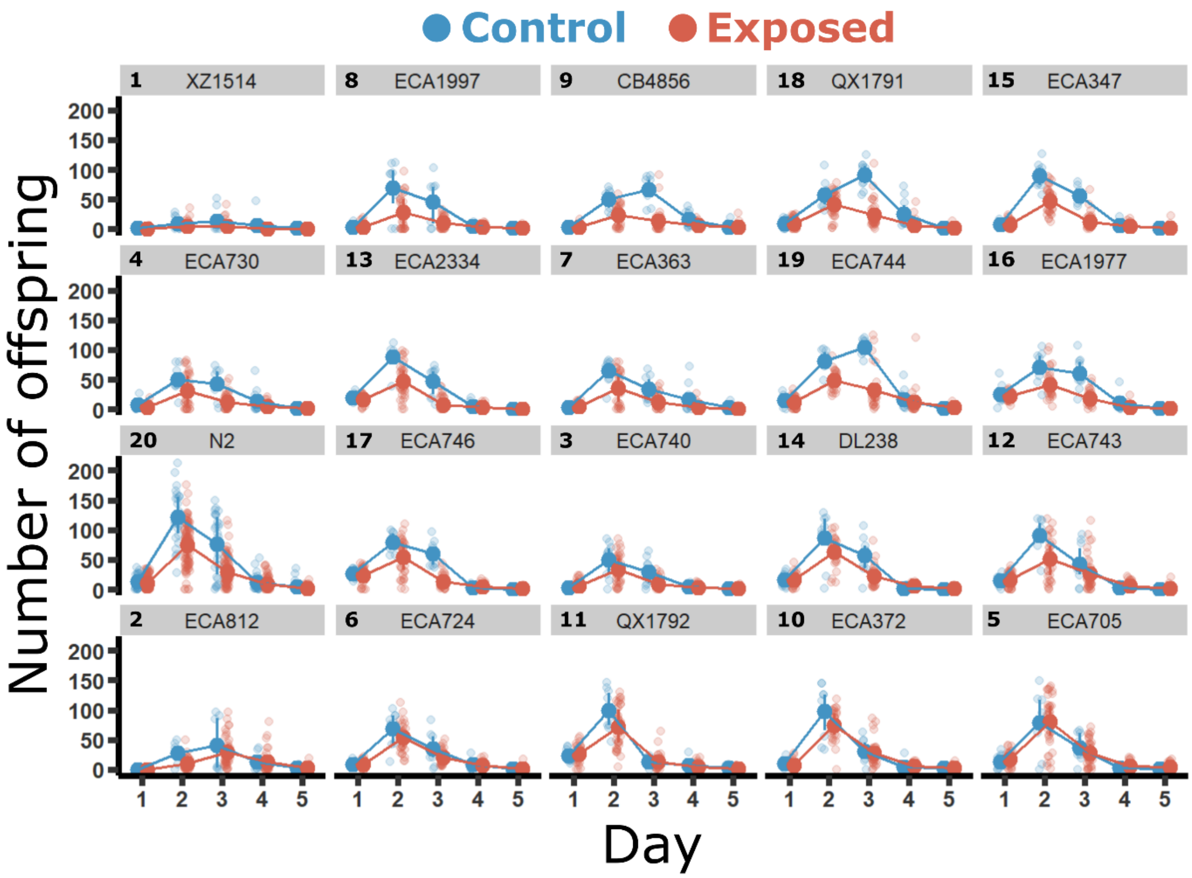


**S4 Fig: Reproductive schedules by host strain and exposure.** As in Fig. 4a,b, solid points indicate the mean number of offspring per host per day, and shaded points show raw data for individual hermaphrodites. Error bars show the interquartile ranges of the data. Blue denotes hosts in the control treatment, and red denotes hosts in exposed conditions, representing all four treatments. Host strains are arrayed from top left to bottom right in order of increasing overall defense against *Nematocida*, as shown in Fig. 2a. Host strains are indicated by both their strain name and numeric ID.

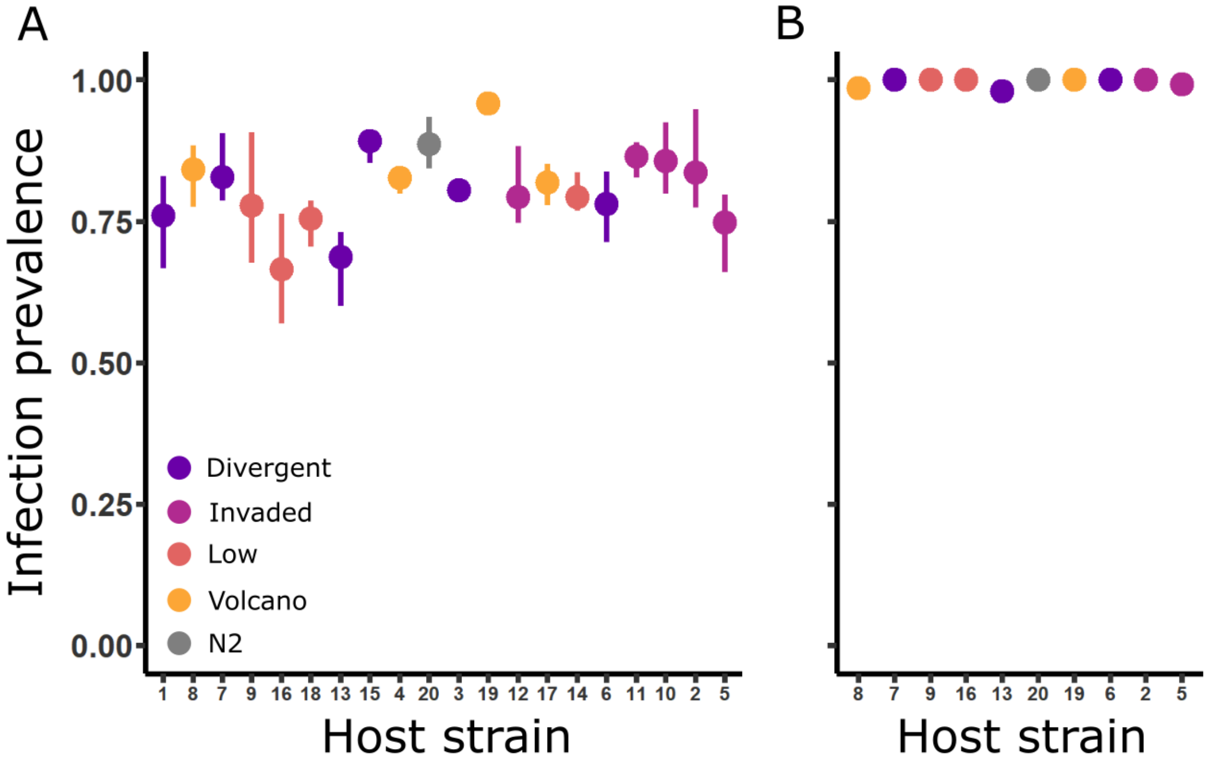


**S5 Fig: Infection prevalence by host strain** at (**A**) 48 and (**B**) 72 hours. Circles indicate the mean prevalence of infection across replicates, and error bars show interquartile ranges of the data. Host strains are arrayed along the x-axis in order of increasing defense against a low dose of *N. parisii*, as shown in Fig. 3b, top left, and they are colored according to host group, with the host strain N2 in gray. Host strains are indicated by their numeric ID.

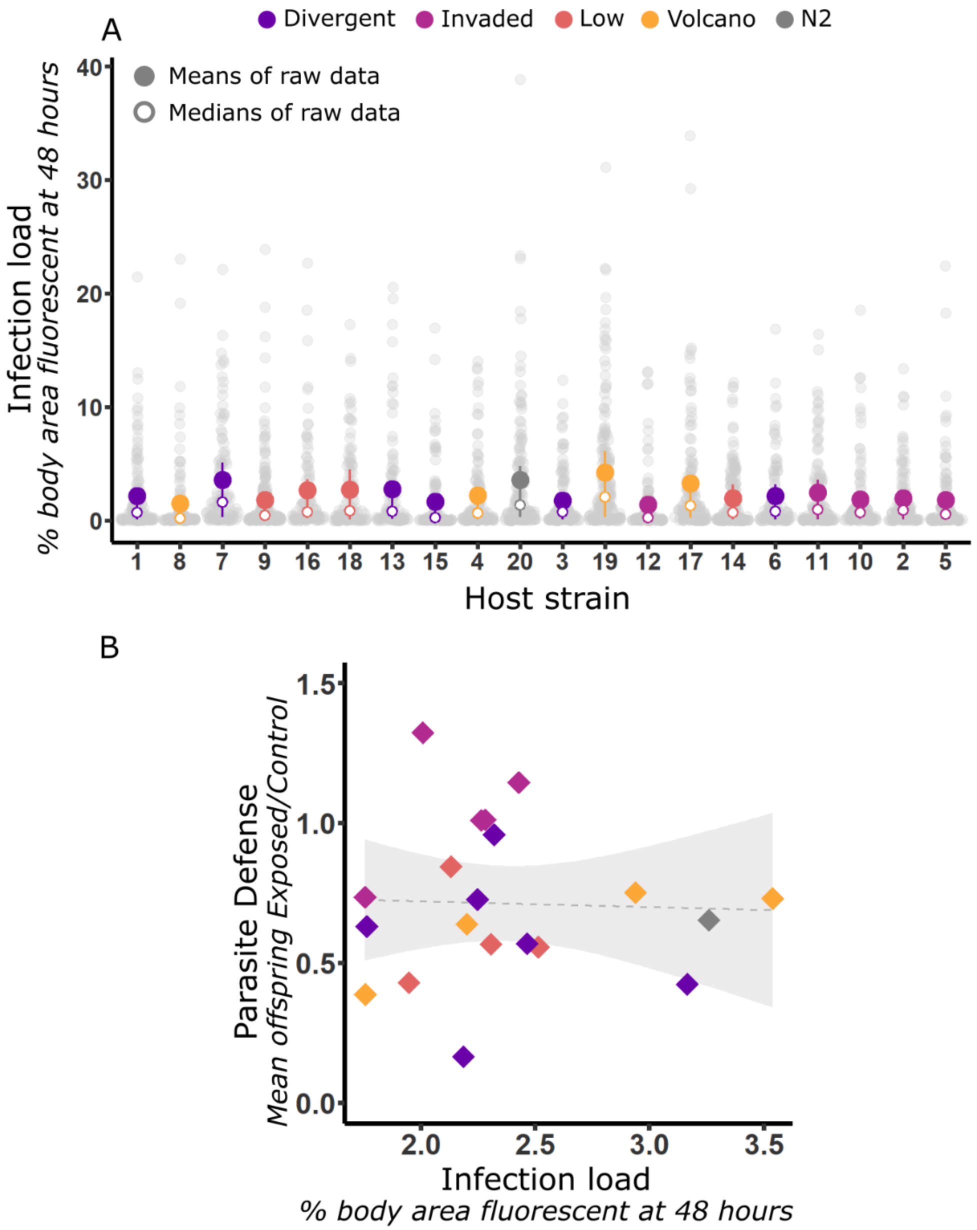


**S6 Fig: Infection load at 48 hours and defense against parasites. A** presents infection load at 48 hours, measured as the percent of body area fluorescent in 2D images of hosts. Higher infection load is interpreted as lower resistance. Gray dots show infection load of individual hosts. Filled circles show means of the raw data plus interquartile ranges. Open circles show medians of the raw data. Host strains are arrayed along the x-axis in order of increasing defense against a low dose of *N. parisii*, as shown in Fig. 3b, top left, and they are colored according to host group, with the host strain N2 in gray. Host strains are indicated by their numeric ID. **B** shows the relationship between infection load at 48 hours and defense. Increasing values on the x-axis indicate decreasing resistance. Each point represents a host strain, colored by host group. Defense is given for the response to a low dose of *N. parisii*, as in Fig. 3b, top left. For load, diamonds show the means of model predictions for replicates from the conditional model in Table P in S1 Text.

**LIST OF WORKS CITED**


1. Troemel ER, Félix M-A, Whiteman NK, Barrière A, Ausubel FM. Microsporidia are natural intracellular parasites of the nematode *Caenorhabditis elegans*. PLoS Biol. 2008;6: e309.

2. Bubrig LT, Janisch AN, Tillet EM, Gibson AK. Contrasting parasite-mediated reductions in fitness within versus between patches of a nematode host. Evolution. 2022;76: 1556–1564. doi:10.1111/evo.14521

3. Therneau T. A Package for Survival Analysis in R. 2024. Available: https://CRAN.R-project.org/package=survival
